# Chromatin accessibility is a two-tier process regulated by transcription factor pioneering and enhancer activation

**DOI:** 10.1101/2022.12.20.520743

**Authors:** Kaelan J. Brennan, Melanie Weilert, Sabrina Krueger, Anusri Pampari, Hsiao-Yun Liu, Ally W.H. Yang, Timothy R. Hughes, Christine A. Rushlow, Anshul Kundaje, Julia Zeitlinger

## Abstract

Chromatin accessibility is integral to the process by which transcription factors (TFs) read out cis-regulatory DNA sequences, but it is difficult to differentiate between TFs that drive accessibility and those that do not. Deep learning models that learn complex sequence rules provide an unprecedented opportunity to dissect this problem. Using zygotic genome activation in the *Drosophila* embryo as a model, we generated high-resolution TF binding and chromatin accessibility data, analyzed the data with interpretable deep learning, and performed genetic experiments for validation. We uncover a clear hierarchical relationship between the pioneer TF Zelda and the TFs involved in axis patterning. Zelda consistently pioneers chromatin accessibility proportional to motif affinity, while patterning TFs augment chromatin accessibility in sequence contexts in which they mediate enhancer activation. We conclude that chromatin accessibility occurs in two phases: one through pioneering, which makes enhancers accessible but not necessarily active, and a second when the correct combination of transcription factors leads to enhancer activation.

## Introduction

Cellular transitions during embryonic development are driven by cis-regulatory DNA sequences, or enhancers, that instruct genes to become expressed at the right time and place. Each enhancer contains a distinct combination and arrangement of sequence recognition motifs for transcription factors (TFs) such that only a specific combination of TFs, present at the right time and place in development, can stimulate activation^1,2^. How exactly combinations of TFs read out the cis-regulatory code to foment enhancer activation is a fundamental question in biology.

An important layer of the cis-regulatory code is chromatin accessibility^3^. Chromatin accessibility both informs and is impacted by the binding of TFs and thus is an integral part of the process by which enhancers become activated. Before activation, developmental enhancers are maintained in a state of intrinsically high nucleosome occupancy such that they are inaccessible to most TFs^4–8^. The first step towards activation is to make the enhancer accessible, which is accomplished by the so-called “pioneer” TFs. Pioneer TFs are typically expressed early during cellular transitions and can bind their motifs within nucleosomal DNA^9–11^. Once the chromatin is accessible, additional TFs may bind to and activate enhancers, leading to the expression of target genes. However, TFs frequently cooperate in modulating chromatin accessibility^12–15^, making it hard to differentiate between pioneer TFs and non-pioneer TFs, and raising the possibility that any TF may function as a pioneer TF^16–18^.

Distinguishing between motifs of TFs that actively drive chromatin accessibility and those of TFs that follow it more passively is computationally challenging. A motif may be statistically overrepresented in accessible regions, but whether it facilitates chromatin accessibility or is present in these regions and subsequently contributes to enhancer activation once the region is already accessible is not clear. Identifying pioneer TFs experimentally is also challenging. In *in vitro* experiments, pioneer TFs have an affinity for nucleosomes and tend to be structurally capable of binding their motif on nucleosomal DNA^19–22^. Thus, pioneers may read out nucleosomal DNA sequences differently than when binding to naked DNA^19,22,23^, but the general rules of these interactions are unknown.

To distinguish pioneer TFs from non-pioneer TFs, one possibility is to model chromatin accessibility data in a high-resolution and quantitative fashion, while taking motif combinations and arrangements into account^18^. This approach is even more powerful when combined with interpretable convolutional neural networks (CNNs), which can learn complex DNA sequence rules embedded in the cis-regulatory code *de novo^24^*. In this learning paradigm, the CNN learns to predict the experimental data directly from genomic sequence, which allows it to learn motifs in their combinatorial context. These rules are general since the performance is evaluated based on a withheld subset of the data that the model does not train on. If the model can accurately predict these test data, the learned sequence rules are extracted from the model using interpretation tools^25^.

This approach has been successfully used to predict ATAC-seq chromatin accessibility data^26–30^, revealing TF motifs predicted to contribute to chromatin accessibility in different experimental systems. However, since not all TFs and their binding motifs are known under these conditions, it is very difficult to evaluate whether the discovered motifs belong to known TFs with characterized properties^31^. Likewise, the models can predict synergistic effects between TF motifs^28,29^, but the exact rules and the underlying mechanisms are not known. This makes it very challenging to connect the rules extracted from deep learning models with known TF biology.

To better leverage this approach, we set out to learn both TF binding data and chromatin accessibility data in the early *Drosophila* embryo, a well-studied model system with a wide range of data from classical genetics, biochemistry, and modern imaging experiments. Studying early embryogenesis has the added advantage that chromatin accessibility is established *de novo* as the zygotic genome is activated and the first gene expression programs are established along the anteroposterior and dorsoventral axes^32–34^. The TFs and enhancers involved in this process have been thoroughly characterized by molecular genetics^35^, making it an ideal system to test and validate the learned rules of a CNN model.

The major driver of the *Drosophila* zygotic genome activation is the maternally-provided zinc-finger TF Zelda, which begins to bind one hour into development, during the embryo’s eighth nuclear cycle^36,37^. From then on, Zelda binds the majority of its motifs genome-wide, which are highly enriched among developmental enhancers^36,38,39^. At these regions, Zelda binding is required for nucleosome depletion and increased chromatin accessibility^6,40,41^. This in turn facilitates the binding of patterning TFs, including the binding of the dorsoventral patterning TFs Dorsal^42,43^ and Twist^44^, as well as the anteroposterior patterning TFs Bicoid^45–47^ and Caudal^5^. Furthermore, *in vitro* experiments suggest that Zelda can bind in the presence of nucleosomes^19,48^. Taken together, Zelda has all the characteristics of a pioneer TF.

While Zelda is a well-studied pioneer TF, whether it cooperates with other early-acting TFs in the embryo to induce chromatin accessibility is not known. GAGA Factor (GAF) and CLAMP are additional pioneer TFs important for zygotic genome activation, but whether they synergize with Zelda is not clear, because they regulate largely distinct sets of regions from Zelda and tend to be more promoter-specific^49–53^. Patterning TFs, on the other hand, strongly overlap in binding with Zelda, but it is unknown whether they cooperate with Zelda and can function as pioneer TFs^36,38,39,54,55^. Bicoid has been reported to play a pioneering role at a subset of its bound regions^56^, but the sequence rules underlying this behavior have not been characterized. Likewise, whether other patterning TFs can increase chromatin accessibility is unknown.

To learn DNA sequence rules at the highest possible resolution, we have previously developed a CNN called BPNet and applied it to high-resolution chromatin immunoprecipitation (ChIP-nexus) data in mouse embryonic stem cells^57,58^. BPNet directly predicts genomics data at base-resolution, allowing it to learn the precise rules by which TFs cooperate in binding *in vivo*. A modified BPNet model, ChromBPNet, has been applied to predict ATAC-seq data at base-resolution^28^, allowing us to use the BPNet approach for both data types.

We generated high-resolution TF binding data and timecourse chromatin accessibility measurements in the early *Drosophila* embryo and leveraged both the unique strengths of the CNN models and our ability to test and validate the learned rules experimentally. We uncovered a clear directional relationship in binding between Zelda and the patterning TFs and found that Zelda and the patterning TFs both increase chromatin accessibility. Through genetic experiments in *Drosophila* mutant strains, we found that Zelda and the patterning TFs increase accessibility through distinct modes. While Zelda acts as a *bona fide* pioneer TF, even at low-affinity motifs, the patterning TFs increase accessibility through transactivation. These results show that chromatin accessibility during zygotic genome activation follows complex sequence rules and is driven both by pioneers and transcriptional activators in distinct steps.

## Results

### Neural networks predict Zelda’s role in helping other transcription factors bind in the early *Drosophila* embryo

To determine the binding and cooperativity of TFs in the early embryo, we performed high-resolution ChIP-nexus experiments in staged embryos on the most well-studied TFs during early embryogenesis. We chose the two best known pioneers, Zelda and GAF, the main dorsoventral patterning TFs Dorsal (Dl) and Twist (Twi), as well as the main anteroposterior patterning TFs Bicoid (Bcd) and Caudal (Cad) (Figure 1a). ChIP-nexus maps genome-wide TF binding footprints at base-resolution by virtue of a strand-specific exonuclease, and has previously uncovered TF cooperativity *in vivo*^57–59^. Replicates for each TF showed high concordance (Supplemental figure 1).

**Figure 1.**
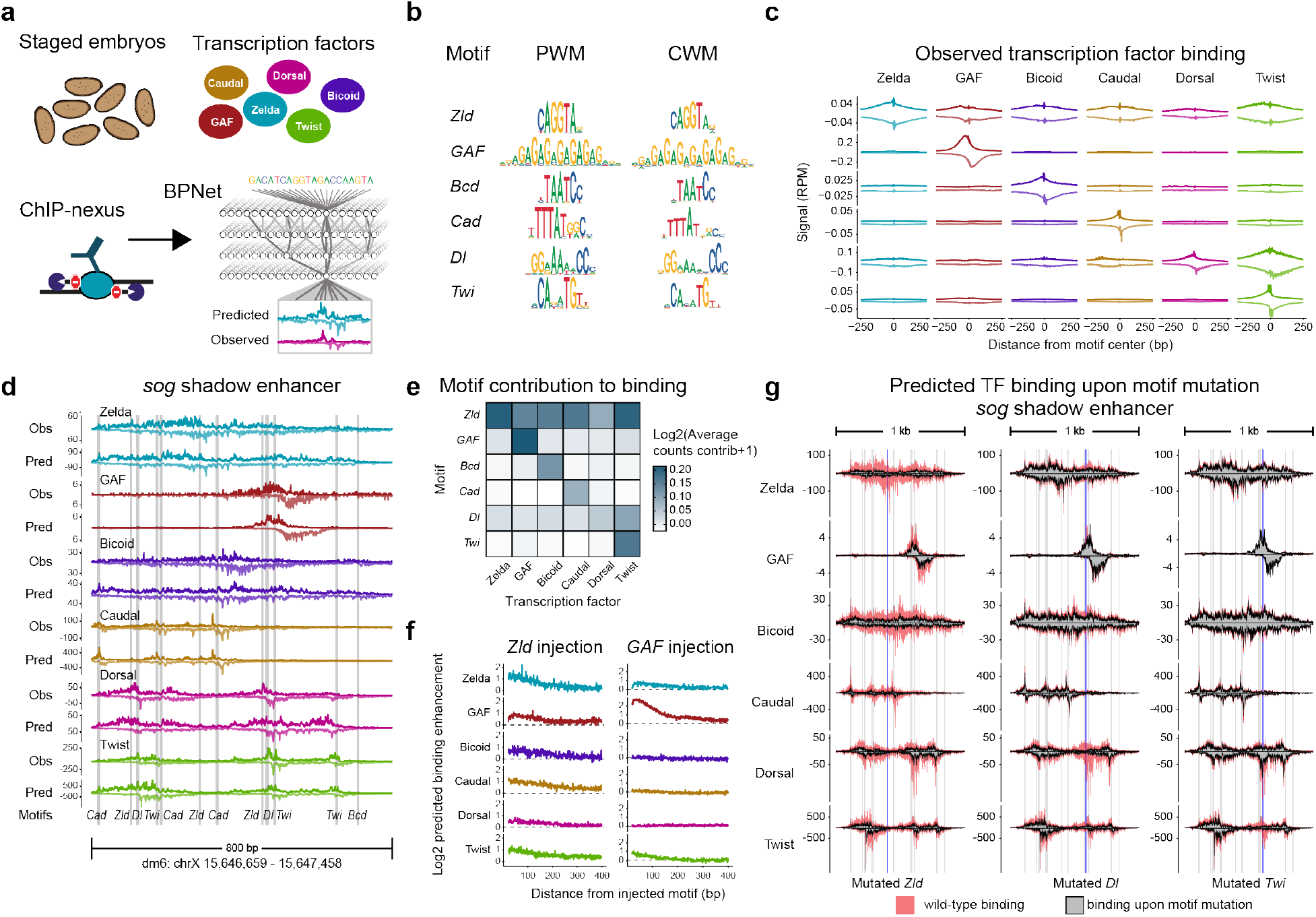
BPNet predicts a hierarchical relationship between Zelda and patterning TFs in the early *Drosophila* embryo. **(a)** Schematic summary of the experimental design. ChIP-nexus was used to map the high-resolution, strand-specific binding of Zelda (Zld), GAGA factor (GAF), Bicoid (Bcd), Caudal (Cad), Dorsal (Dl), and Twist (Twi) in staged syncytial blastoderm embryos. These data were used to train a multi-task BPNet model that predicts TF binding from DNA sequence alone. **(b)** BPNet identified and mapped the known motifs for each TF. The position weight matrix (PWM) is a frequency-based motif representation, while the contribution weight matrix (CWM) is the novel BPNet motif representation, where base height reflects the importance for predicting TF binding. PWM and CWM motif representations are highly similar for all TFs. **(c)** Average TF binding footprints at all BPNet-mapped motifs from the experimentally generated ChIP-nexus data. Sharp binding footprints indicate that a motif is directly bound by a particular TF. Profiles are centered on motifs and binding signals are normalized (RPM). ChIP-nexus provides strand-specific information, with the positive strand represented by positive values and the negative strand represented by negative values. **(d)** Comparing experimentally generated TF binding with BPNet-predicted TF binding at the *sog* shadow enhancer illustrates BPNet’s predictive accuracy. Each color is a different TF, where the top track is the experimental ChIP-nexus data, and the bottom track is the predicted binding. Motifs were identified and mapped by BPNet. This enhancer was withheld from BPNet during training, making it an ideal locus to test how well BPNet has learned the cis-regulatory rules that predict TF binding. **(e)** The counts contribution score for each motif was calculated and averaged for all mapped motifs for the binding of each TF. Darker colors indicate that a motif (y-axis) has a higher contribution to the binding of the associated TF (x-axis). The Zelda motif has a high contribution for the binding of all TFs, but not the reverse, indicating a hierarchical relationship. **(f)** The Zelda motif is predicted to boost the binding of all TFs, while the GAF motif boosts only GAF’s binding. All TF motifs were injected into randomized sequences and their binding was predicted by BPNet when each motif was alone and when a Zelda motif was injected at a given distance, up to 400 bp away. The same procedure was repeated by injecting GAF motifs at distances up to 400 bp away from all other injected motifs. Fold-change binding enhancements were calculated from predicted TF binding in the presence of Zelda/GAF and when these motifs weren’t injected for every distance between motifs (x-axis). **(g)** BPNet predicts TF binding at the wildtype *(wt)* sequence of the *sog* shadow enhancer and when individual motifs are computationally mutated. Shaded colors represent the *wt* predicted binding for each of the six TFs across the entire enhancer. Gray-filled profiles represent the predicted TF binding in response to mutating either a Zelda motif (left), Dorsal motif (middle), or Twist motif (right). Blue bars highlight the mutated motifs in each of the three predictions, while gray bars are all other mapped motifs across the enhancer. Mapped motifs are the same as those highlighted in Figure 1d. Mutating the Zelda motif reduced all TF binding across the enhancer, while mutating the Dorsal motif had a smaller but notable effect on TF binding.

We trained a BPNet model to predict the ChIP-nexus data from DNA sequence and interpreted the sequence rules as previously described^57^. This approach is uniquely suited to learn the sequence rules of TF binding and cooperativity because it models cis-regulatory sequences in their native genomic contexts and learns TF binding motifs in an inherently combinatorial way. Motifs that are mapped in genomic sequences are defined not just by a sequence match but also by a contribution score towards the binding predictions. To maximize the accuracy of the model’s learned sequence rules, we optimized the model to achieve high prediction accuracy and confirmed the results through crossvalidation (Supplemental figure 2).

We next inspected the *de novo* learned motifs, represented either as a classic frequency-based position weight matrix (PWM) or as the novel contribution weight matrix (CWM), which is the model’s extracted contribution of each base for TF binding. This confirmed that we discovered the known motifs for all BPNet-modeled TFs (Figure 1b) and that these motifs showed the expected sharp ChIP-nexus binding footprints from the bound TFs (Figure 1c). We also manually inspected well-studied enhancers to compare how the ChIP-nexus predictions matched the experimental data and that experimentally validated motifs were mapped accurately (Figure 1d, Supplemental figure 3). For example, we confirmed that the well-studied neuroectodermal *sog* shadow enhancer had the expected motifs for Zelda and Dorsal^42,43,60,61^ and for Twist and Bicoid^62–64^. Since this enhancer is part of the withheld data set that was never seen by the model during training, this example highlights how the model correctly predicts TF binding from DNA sequence alone and that it did so by using the expected TF binding motifs (Figure 1d).

We then extracted the rules of TF cooperativity from the model. We first measured the average contribution of each motif towards the binding of each TF (Figure 1e). As expected, all motifs strongly contributed towards their own TFs, but some motifs also contributed to the binding strength of other TFs, suggesting that there is binding cooperativity between TFs. Most prominently, the Zelda motif is predicted to be important for the binding of all other TFs (Figure 1e). This includes Bicoid, Caudal, Dorsal, and Twist, which have been shown in previous genetic experiments to depend on Zelda binding to its motif and thus agrees with Zelda’s established role as a pioneer TF^5,6,40,42,44,45^. In addition, BPNet predicts that Twist binding depends on the Dorsal motif. Dorsal and Twist have previously been reported to cooperate^61,65–68^, but our result suggests that this cooperativity is directional, i.e., the Dorsal motif is more important for Twist binding than the Twist motif is important for Dorsal binding. This is also reflected in the experimental ChIP-nexus average profiles, which show Twist accumulation over the Dorsal motif but not vice versa (Figure 1c). Interestingly, the motif for GAF did not strongly contribute to the binding of TFs other than GAF itself, even though GAF is known to promote chromatin accessibility^49,52,53,69,70^.

To internally validate that BPNet learned different rules of cooperativity for Zelda and GAF, we used the trained model to predict TF binding when motif pairs are injected into randomized sequences (Figure 1f). For each TF motif, we measured the average fold-change increase in binding when a Zelda or GAF motif was added at a given distance (up to 400 bp). Consistent with our initial results, injecting a Zelda motif generally boosted the binding of all TFs, while the GAF motif only had a strong boosting effect on another GAF motif (Figure 1f). Notably, all observed cooperativity occurred when the motifs were spaced within nucleosome-range distances, consistent with an effect on nucleosomes.

Finally, we tested the derived cooperativity rules on known enhancers. We computationally mutated the sequence of each TF motif and predicted the effects on TF binding with BPNet. As expected, mutating Zelda motifs consistently had a strong effect on the binding of other TFs (Figure 1g; Supplemental figure 4). In contrast, the effects of mutating patterning TF motifs tended to be more enhancer-specific. At the *dpp* enhancer, mutating Dorsal motifs affected Dorsal and Twist binding, as expected (Supplemental figure 4). However, at the *sog* shadow enhancer, mutating a Dorsal motif also had an effect on the binding of other TFs, including Bicoid (Figure 1g). Likewise, mutating a Twist motif not only affected Twist binding, but also had a weak effect on Dorsal binding. These results suggest more complex rules at some enhancers and raise the question of whether chromatin accessibility plays a role in the observed cooperativity.

### The sequence rules for chromatin accessibility reveal motif-driven pioneer transcription factors

To understand the relationship between TF binding and chromatin accessibility, we performed ATAC-seq experiments^71,72^ in a developmental time course of 30-minute intervals during the maternal-to-zygotic transition. This allowed us to measure how enhancers transition over time from a naturally closed state to a more accessible, primed state^50,73–77^. The first embryo collection (1-1.5 h after egg laying, AEL) covers the time when Zelda begins to bind throughout the genome in the 8th nuclear cycle^36^, as well as the earliest stages of embryonic patterning. During the second collection (1.5-2 h AEL), patterning TFs become active and the major burst of zygotic transcription begins and continues into the third (2-2.5 h AEL) and fourth (2.5-3 h AEL) collections^34,78^. All experiments were performed in triplicate, with highly correlated replicates (Supplemental figure 5). In agreement with previous studies, we find that genome-wide chromatin accessibility increases over the four time points^50^ (Figure 2a).

**Figure 2.**
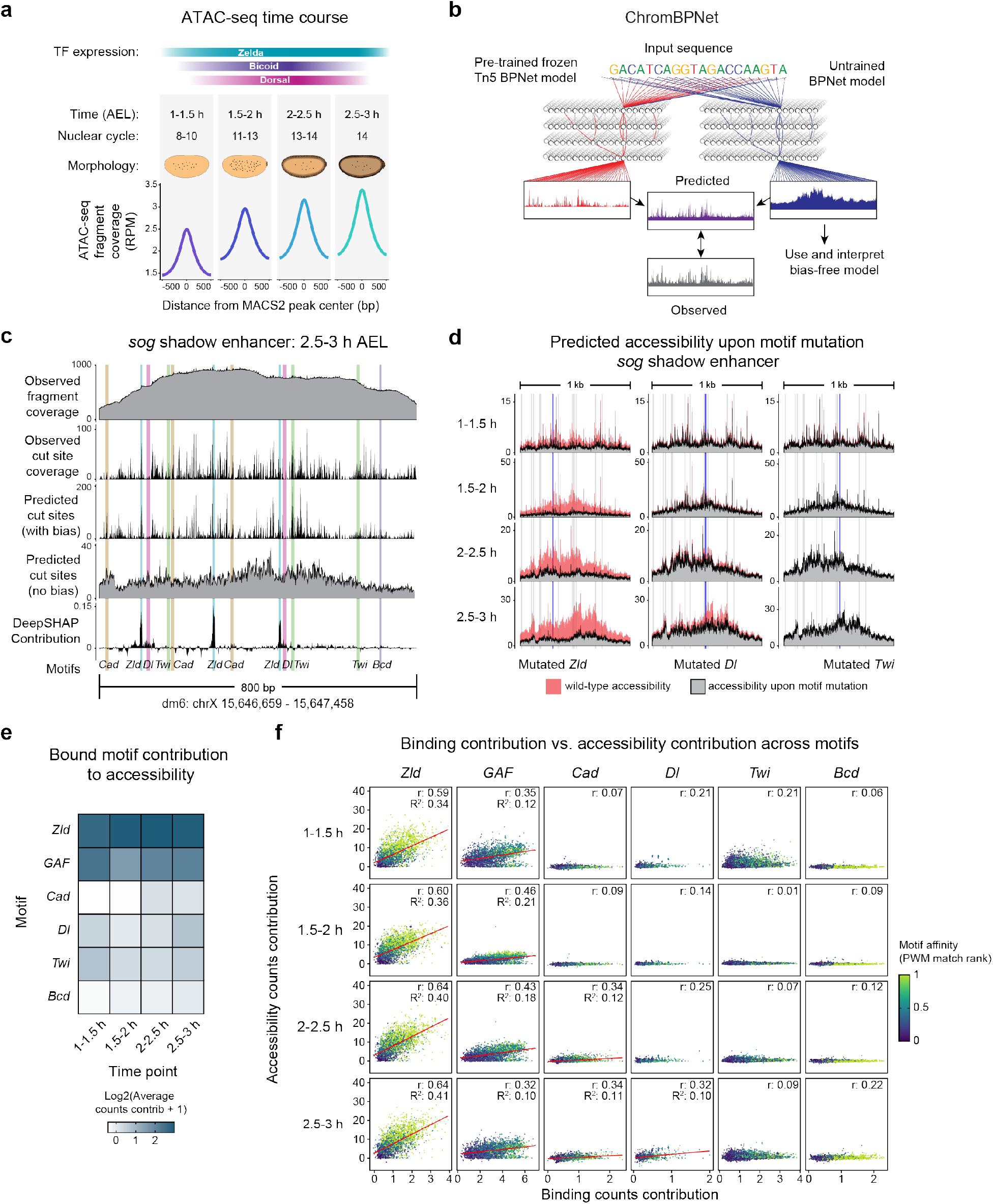
ChromBPNet reveals distinct contribution from pioneers and patterning TFs in early *Drosophila* embryos. **(a)** Schematic summary of ATAC-seq time course experiments. ATAC-seq experiments were performed in four 30-minute windows during the *Drosophila* maternal-to-zygotic transition. Zelda binding begins during the first time point, and patterning TFs start binding in the second time point and increase for the remaining time points. These windows are characterized by distinct embryo nuclear division cycles and morphological features, which facilitated the precise hand-sorting of different stages during embryo collections. Across ATAC-seq peaks there is a general increase in normalized ATAC-seq fragment coverage over time. **(b)** ChromBPNet is a modified BPNet deep learning model that predicts chromatin accessibility using DNA sequence as an input. ChromBPNet’s architecture is similar to the BPNet architecture, however training relies on the simultaneous use of two models. The first is a Tn5 bias model, which was pre-trained on closed and unbound genomic regions to explicitly learn only Tn5 sequence bias and is then frozen. The second is a standard, randomly-initialized BPNet model which learns the unbiased cis-regulatory information predictive for chromatin accessibility. Following model training, the Tn5 bias model is removed, and the unbiased model is interpreted free of Tn5 bias. **(c)** ChromBPNet accurately predicts chromatin accessibility information at the *sog* shadow enhancer during the last time point. The top two tracks represent experimentally generated ATAC-seq coverage, with the top being the conventional fragment coverage and the bottom being Tn5 cut site coverage. The third track is ChromBPNet’s ATAC-seq cut site prediction at this time point. While it mirrors the observed cut site coverage very closely, this track contains Tn5 bias. The fourth track is ChromBPNet’s prediction after removing Tn5 bias, which is more evenly distributed across the enhancer. The fifth track is the counts contribution scores for each base across the enhancer, which spikes at BPNet-mapped motifs, particularly at Zelda motifs but also at Dorsal motifs. **(d)** ChromBPNet predicts chromatin accessibility at the wildtype *(wt) sog* shadow enhancer and in the presence of individual motif mutations across time. The same Zelda (left), Dorsal (middle), and Twist (right) motifs that were mutated previously (Figure 1g) are mutated here, and ChromBPNet predicted time course chromatin accessibility in response to those mutations. Mutation of the Zelda motif had the largest predicted effect on chromatin accessibility, while the Dorsal mutation is predicted to lower accessibility to a lesser extent and only at later time points. Shaded colors are the *wt* predicted accessibility for each time point, and the gray profiles are the predictions in response to motif mutation. Blue bars are mutated motifs, gray bars are all other motifs mapped to this enhancer. **(e)** Average counts contribution scores for each BPNet-mapped motif (y-axis) are shown for all time points (x-axis) to represent how important a particular motif is to the chromatin accessibility prediction across time. Pioneering motifs contribute to chromatin accessibility robustly at all time points, while patterning TF motifs have a lesser contribution that is limited to later time points. **(f)** Pioneer TF motifs show a clear three-way correlation between binding contribution, accessibility contribution, and motif strength. Patterning TFs show much weaker, time point-specific relationships, suggesting context-dependent behavior. For each bound and accessible motif for all TFs, the binding counts contribution scores (x-axis) and accessibility counts contribution scores (y-axis) are plotted. Motif strength was extracted from the trained BPNet model by ranking motifs for each TF by their match score to each TF’s PWM and taking the rank percentile. Pearson correlation values (r) and coefficient of determination R^2^ values were calculated. Red lines are shown for plots with an r > 0.3.

In order to understand the cis-regulatory sequence rules that guide these chromatin accessibility data, we used ChromBPNet, a variation of BPNet that predicts ATAC-seq data at the highest resolution^28,79^. Rather than training on whole fragment coverage, the model predicts the cut sites made by the Tn5 transposase, which more accurately represent accessibility (Figure 2b). Since the Tn5 transposase possesses a strong sequence bias in its cut position^80,81^, ChromBPNet is designed to remove this experimental bias by explicitly learning its sequence rules in a separate BPNet model trained on closed genomic regions (i.e., with low-count, non-peak ATAC-seq signal) (Figure 2b). Then a second ChromBPNet model learns how sequence influences the ATAC-seq accessible regions beyond the bias that is already captured by the frozen bias model (Supplemental figure 6a-b). After training, the bias model is removed, and the second model is interpreted to extract the biologically relevant sequence rules that predict chromatin accessibility.

We trained separate ChromBPNet models for each of the ATAC-seq time points, omitting regions with annotated promoters to ensure that the sequence rules learned were specific for enhancers, and not strongly driven by core promoter motifs. As with BPNet, we computed performance metrics, conducted hyperparameter tuning, and trained crossvalidation models to validate that model training was successful (Supplemental figure 6c-e).

To visually inspect ChromBPNet’s predictions, we used the *sog* shadow enhancer as example (Figure 2c; additional enhancers in Supplemental figure 7). The observed cut site coverage from the ATAC-seq data were spiky and without discernible footprints around the known motifs. This pattern closely matched the cut site coverage predicted by the model, consistent with its high performance metrics (Supplemental figure 6c-e). After removing the bias, the predicted chromatin accessibility was more evenly distributed over the entire enhancer, suggesting that the Tn5 cut site bias was successfully removed (Figure 2c).

As with BPNet, we extracted base-resolution contribution scores for all sequences and summarized the *de novo* learned motifs. The motifs for Zelda and GAF were robustly re-discovered at all four time points, consistent with them being pioneer TFs that open chromatin (Supplemental figure 6f). Additionally, the accessibility model discovered Caudal-like, Dorsal-like, and Twist-like motifs, which deviated from those learned by the TF binding model but nevertheless showed the expected ChIP-nexus binding footprints, confirming their identity (Supplemental figure 6f). We did not identify the Bicoid motif, which seems to contradict a previous study suggesting a role for Bicoid in chromatin accessibility, however this role was context-dependent, and thus the underlying sequence rules were unclear^56^.

We next critically evaluated whether the learned sequence rules were compatible with previous knowledge from known enhancers. When we inspected the contribution scores, we found that the Zelda motifs typically stood out with high scores, but some Dorsal and Caudal motifs also had contribution, confirming that these motifs were learned (Figure 2c, Supplemental figure 7). As another way of internally validating the importance of these motifs, we performed *in silico* mutagenesis (Figure 2d; Supplemental figure 8). As expected, mutating a Zelda motif in the *sog* shadow enhancer strongly reduced the predicted chromatin accessibility for all time points, but mutating a Dorsal motif also weakly reduced the predicted accessibility, especially at the later time points when patterning TFs bind most strongly^5,78^. Taken together, the interpretations agree with the TF binding model and our understanding of patterning TF binding dynamics in the *Drosophila* embryo, suggesting that it is a useful model to probe the rules of chromatin accessibility.

We next set out to systematically compare the rules of binding with those of accessibility. We selected regions that are accessible and contain TF motifs mapped by the binding model, which ensures that the motifs are high-quality and unambiguously mapped to the TF through a direct sequence-to-binding relationship. We confirmed that the Zelda and GAF motif instances had high contribution to accessibility at all time points, while those of the patterning TFs had a much smaller contribution (Figure 2e). Similar effects were observed when we injected each TF motif into randomized sequences *in silico* (Supplemental figure 6g). Using these mapped motif instances, we then plotted the predicted contribution to accessibility as a function of the predicted binding contribution (Figures 2f).

If the role of patterning TFs is more context-dependent than *bona fide* pioneer TFs, we would expect pioneer TFs to have a more consistent relationship between the TF’s binding and the generated chromatin accessibility. Indeed, a correlation between total Zelda binding and chromatin accessibility has previously been reported^40,42^, but it is unknown how well this holds for individual motifs and how this compares to other TFs. Strikingly, we observed a strong correlation for both Zelda and GAF motifs between accessibility and binding contributions, in spite of being learned by different models on different types of data (Figure 2f). Moreover, when we derive a simple score for motif strength (rank percentile of the PWM match scores), we see that binding and accessibility contributions increase as motif strength increases. This three-way association suggests that the accessibility generated by Zelda and GAF is motif-driven and not heavily reliant on the surrounding enhancer context, which agrees with the conventional model that pioneer TFs come first and mediate the initial step in enhancer activation.

In contrast, when we plot the same correlations for the patterning TFs, we find much weaker relationships between TF binding and chromatin accessibility (Figure 2f). Here, stronger measures of motif strength are associated with stronger binding contribution but not accessibility contribution. One exception is Dorsal at the last time point, where we find an increased correlation between binding and accessibility contribution (Pearson correlation 0.32), as well as an association with motif strength. Notably, this occurs when Dorsal’s binding has been reported to be strongest during development^5^. Likewise, Caudal also has a time point-specific correlation that is highest at the two latest time points, when its binding is also strongest^5^. For Twist and Bicoid motifs, the binding and accessibility contribution correlation is the poorest, consistent with the difficulty of the model discovering their canonical motif representations. Taken together, our binding and accessibility models suggest an operational definition of pioneer TFs in which pioneer TFs open chromatin in a motif-driven fashion, while other TFs may also play a role in increasing chromatin accessibility but do so in a more context-dependent manner.

### Zelda’s effect on opening chromatin extends to low-affinity motifs

The correlation between motif strength, TF binding, and ability to open chromatin implies that motifs of lower affinity can also pioneer chromatin accessibility but do so proportionally less than high-affinity motifs. This is surprising since pioneering is expected to occur through TF binding on nucleosomes, where sequence recognition is structurally more constrained than on naked DNA^10,19,20,82–84^. Given previous evidence that pioneering events identified *in vivo* were associated with degenerate motifs^22,23^, we set out to validate the prediction that pioneering by Zelda can involve low-affinity motifs.

We first examined whether the BPNet models correctly learned motif affinities from the Zelda ChIP-nexus binding data. We took all bound Zelda motifs mapped by BPNet and plotted their sequences ordered by contribution to Zelda binding (Figure 3a). The motif that contributed most to binding (sequence logo from the top quartile) was the canonical CAGGTAG motif, while low-affinity binding motifs (sequence logo from bottom quartile) included motifs where the last base was not a G (CAGGTAH), or the first base was a T (TAGGTAG). These results are consistent with the Zelda motif affinities determined previously by gel shift studies and mutant data^36–38,85,86^ and correlate with the observed chromatin accessibility across these motifs (Supplemental figure 10a).

**Figure 3.**
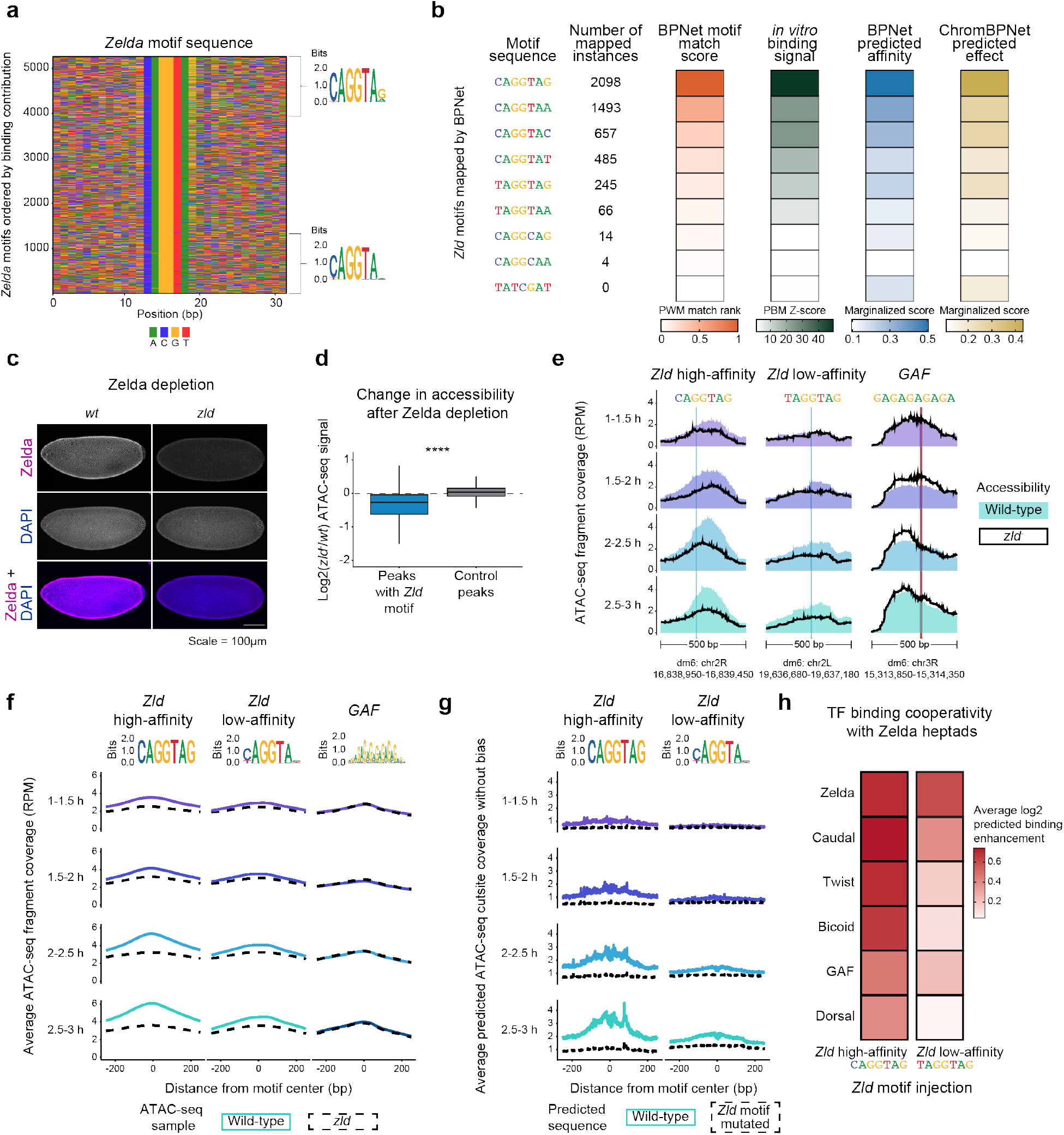
The pioneer TF Zelda reads out motif affinity to drive chromatin accessibility. **(a)** BPNet binding contributions reflect the known Zelda motif affinities. All BPNet-mapped Zelda motifs were ordered by their counts contribution scores to Zelda binding, with the highest contribution motifs on top and the lowest contribution motifs on the bottom. Motif logos were generated for the highest and lowest contributing sequence quartiles. **(b)** Zelda motif affinities can be accurately extracted from the trained BPNet and ChromBPNet models. Mapped Zelda motifs were separated by their heptad sequences, and the known Zelda heptads were extracted and ordered by their rank percentile of their PWM match scores (orange). The negative control, non-mapped TATCGAT heptad was included. Protein binding microarray (PBM) experiments were performed using the Zelda C-terminal region. 8-mers from PBM experiments were grouped based on their 7-mer sequences, median Z-score (green) values were calculated for the 7-mers, and the 7-mers matching Zelda heptads were extracted. The effects of each Zelda heptad were marginalized from the effects of genomic background sequences using the trained BPNet (blue) and ChromBPNet (gold) models to extract model-determined motif affinities. The experimentally derived and model-derived Zelda motif affinities strongly correlate. **(c)** Zelda depleted embryos *(zld^-^)* show a clear reduction in the Zelda protein. Confocal images of nuclear cycle 14 wildtype *(wt)* and *zld^-^* embryos were collected, maximum intensity projected, and processed using the same settings. **(d)** Chromatin accessibility is significantly reduced at ATAC-seq peaks containing mapped Zelda motifs. Differential chromatin accessibility between *wt* and *zld^-^* embryos was calculated as the log2 fold change for each peak region using DESeq2. The median values of the four time points are shown. Peaks containing Zelda motifs are significantly different from control peaks without Zelda motifs (Wilcoxon rank-sum test, p < 2e-16). **(e)** Chromatin accessibility is reduced at high- and low-affinity Zelda motifs in *zld^-^* embryos. Individual examples of normalized chromatin accessibility in *wt* (shaded profile) and *zld^-^* (black line) embryos are shown at a high-affinity Zelda motif (CAGGTAG, left) and a low-affinity Zelda motif (TAGGTAG, middle), with the GAF motif (right) as a control. No other BPNet-mapped motifs are within these windows. **(f)** Average chromatin accessibility profiles at the 250 highest and lowest affinity Zelda motifs in *wt* and *zld^-^* embryos show that low-affinity motifs facilitate Zelda’s pioneering, but to a lesser extent than the high-affinity motifs. Islands that only contain a single Zelda motif were extracted and separated into high- and low-affinity categories based on the rank percentile of their PWM match scores (high = high affinity, low = low affinity), while 250 GAF motifs were seed-controlled, randomly selected. Motif logos were generated from these motif instances. The colored lines are the *wt*, normalized, ATAC-seq data, and dotted black lines are the same but in *zld^-^* embryos, with profiles anchored on the Zelda motifs. Motifs mapping to promoters were excluded, as in ChromBPNet training. **(g)** ChromBPNet model predictions at the same high- and low-affinity Zelda motifs as in Figure 3f. ChromBPNet predicted bias-corrected cut site coverage at the *wt* high- and low-affinity Zelda motif regions and when the Zelda motifs were computationally mutated. The similarity to Figure 3f shows that ChromBPNet has accurately learned the effects of Zelda motif affinity. **(h)** Low-affinity Zelda motifs are predicted to boost TF binding. TF motifs were injected into randomized sequences with either a high-affinity Zelda motif (CAGGTAG), low-affinity Zelda motif (TAGGTAG), or no Zelda motif injected at a given distance away for up to 200 bp, and TF binding was predicted (y-axis). The fold change binding enhancement averaged across the window was calculated using predicted TF binding at motifs with a high- or low-affinity Zelda motif injected nearby and predicted TF binding without a Zelda motif injected nearby.

To more comprehensively test how well the BPNet models learned relative Zelda motif affinities, we performed *in vitro* protein binding microarray (PBM) experiments^87,88^ for Zelda (Figure 3b). PBM-extracted affinities have been shown to correlate with K_d_ affinity measurements^89–91^. We calculated the median Z-score of the binding signal and its corresponding median E-score for all relevant Zelda motif heptads, as well as a negative control sequence (TATCGAT) used previously in gel shift experiments^38^. Strikingly, the simple BPNet-derived motif strength scores we used earlier closely matched the *in vitro* PBM binding signal (Figure 3b; Supplemental figure 10b). For example, both the experimental data and the BPNet-derived motif strength scores showed on average a three-fold difference in affinity between the CAGGTAG and TAGGTAG sequences.

These results are consistent with the recent finding that accurate predictions of relative motif affinities can be extracted from a BPNet model trained on ChIP-nexus or ChIP-seq data^92,93^. Such relative motif affinities can be derived without using their motif representations by simply predicting TF binding on motif instances that are stripped from the surrounding genomic context. To test this, we “marginalized” each Zelda motif by injecting it into randomized sequences and measured the effects on binding and chromatin accessibility. The log-transformed measurements were very similar to our previous BPNet-derived motif strength scores, and closely matched the *in vitro* PBM binding Z-scores (Figure 3b). These results collectively confirm that the models have accurately learned relative Zelda binding affinities.

Having confirmed that the BPNet and ChromBPNet models correctly learned Zelda motif affinities, we next performed experiments on Zelda-depleted embryos^6^ to test whether low-affinity motifs contribute to accessibility *in vivo*. We confirmed that the *zld^-^* embryos had no detectable Zelda by immunostaining (Figure 3c) and performed ATAC-seq time-course experiments, with replicates that were highly correlated (Supplemental figure 9). Consistent with previous observations^40,56^, Zelda-bound regions showed a global decrease in accessibility compared to wildtype (p < 2e-16, Wilcoxon rank-sum test), while regions without a Zelda motif remained unchanged (Figure 3d; Supplemental figure 10c).

We then asked whether individual low-affinity Zelda motifs by themselves influence chromatin accessibility. We selected regions with either a single high-affinity (CAGGTAG) or a single low-affinity (TAGGTAG) Zelda motif, with no other BPNet-mapped motif nearby. At regions with the high-affinity Zelda motif, a clear reduction in chromatin accessibility was observed in *zld^-^* embryos. This reduction became more prominent over time as these regions became more accessible in wildtype embryos (example in Figure 3e, left). At regions with the low-affinity TAGGTAG motifs, we observed the same effect but weaker (example in Figure 3e, middle). To quantify this difference, we selected the genomic regions with the 250 highest and lowest affinity Zelda motifs. To minimize confounding effects, these regions had no other mapped motifs nearby and did not overlap promoters. As expected, the regions with the high-affinity Zelda motifs had more Zelda binding in the ChIP-nexus data than those with the low-affinity motifs (Supplemental figure 10d). Using these regions, we found that the low-affinity Zelda motifs had on average a fivefold weaker effect on chromatin accessibility than the high-affinity Zelda motifs, while control regions with a single GAF motif were unchanged (Figure 3f; Supplemental figure 10e). These differences were very similar to those predicted by the ChromBPNet upon mutating the Zelda motifs (Figure 3g). These results demonstrate that low-affinity Zelda motifs can promote accessibility, but to a lesser extent than high-affinity CAGGTAG motifs, and that the extent of chromatin opening correlates with the motif’s affinity.

Since the low-affinity Zelda motifs have a smaller effect on chromatin accessibility, we expected them to also have a weaker effect on promoting the binding of patterning TFs. To test this hypothesis, we performed *in silico* motif injections and measured the average predicted binding of each TF with and without the presence of different Zelda motif variants. For all TFs, the resulting fold-change binding enhancement was indeed higher for the high-affinity CAGGTAG motif than for the low-affinity TAGGTAG motif, but the latter still had a measurable effect (Figure 3h). Likewise, the accessibility model predicted that both high- and low-affinity Zelda motifs boosted the effect of patterning TF motifs on chromatin accessibility, but to a different extent (Supplemental figure 10f-g). These effects are consistent with the experimentally observed effect of low-affinity motifs on chromatin accessibility and corroborate the role of low-affinity Zelda motifs in opening chromatin and helping patterning TFs bind.

### Patterning transcription factors contribute to chromatin accessibility

Thus far, the results suggest that patterning TFs do not have the same pioneering capabilities as Zelda, but could increase chromatin accessibility in some contexts, perhaps dependent on which other motifs are present within that region. To systematically investigate motif combinations, we used a “motif island” approach in which genomic regions are grouped according to their motif combinations. An island is initially defined as 200 bp centered on a motif, but if this region overlaps with another motif island, the islands get merged (Figure 4a). We then classified the motif islands by their motif combinations without taking motif number or order into account (islands provided in Supplemental file 2). These multi-motif islands are the size of typical enhancers^94^, with the majority of them being between 200 and 300 bp wide (Supplemental figure 12b). To better characterize enhancer states for different motif combinations, we used staged embryos and performed micrococcal nuclease digestion with sequencing (MNase-seq) and ChIP-seq experiments for the histone modifications H3K27ac and H3K4me1, with highly correlated replicates (Supplemental figure 11). We then analyzed the properties of each island combination (Figure 4b, individual examples in Figure 4c).

**Figure 4.**
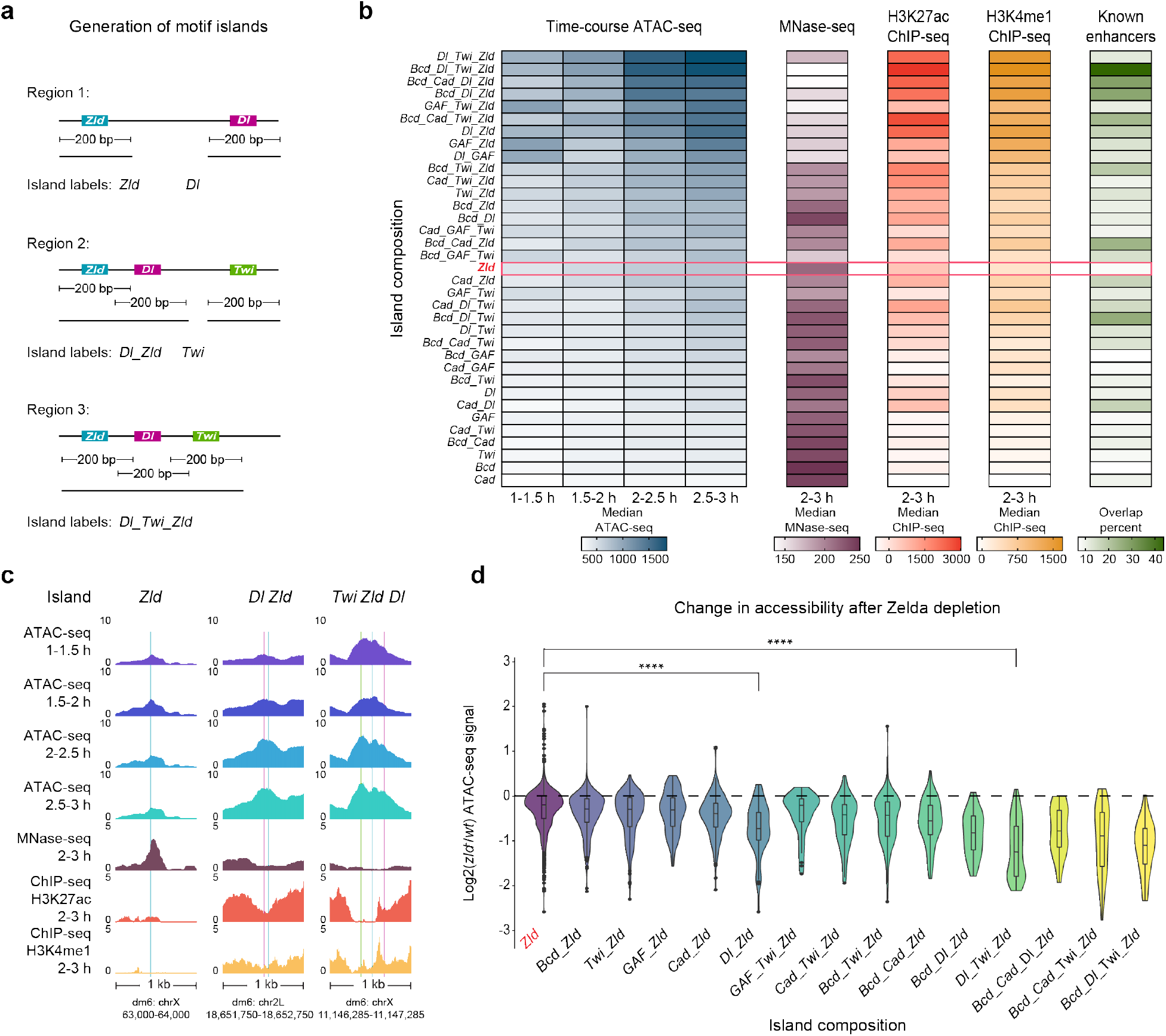
Patterning TFs increase chromatin accessibility in a context-dependent manner. **(a)** Schematic summary of motif islands. Motif islands are generated by first resizing all BPNet-mapped and bound motifs to 200 bp wide. Next, all overlapping regions are reduced together into the final motif islands, and the islands are classified based on the motifs that compose them. This way, all single motif-containing islands (e.g., *zld* only islands) are 200 bp wide. **(b)** Islands with combinations of Zelda and patterning TF motifs contain the highest chromatin accessibility, nucleosome depletion, active enhancer histone modifications, and known enhancer overlap. Motif islands of the same composition are grouped together, and for each island type (y-axis) the median normalized ATAC-seq fragment coverage, MNase-seq signal, H3K27ac ChIP-seq signal, and H3K4me1 ChIP-seq signal were calculated. ATAC-seq and MNase-seq coverage was calculated across a 250 bp window centered on the island, while the H3K27ac and H3K4me1 signals were calculated in a 1.5 kb window centered on the island since these marks are typically on the enhancer flanks. A list of enhancers active in 2-4 h AEL embryos was used to calculate an overlap percentage for each island type^73^. The red bar highlights islands that contain only Zelda motifs, and islands are ordered by total ACAT-seq signal. **(c)** Individual examples for *zld*, *Dl_Zld*, and *Dl_Twi_Zld* islands. Colored bars indicate BPNet-mapped motifs (blue =*zld*, magenta =*Dl*, green =*Twi*), and no other BPNet-mapped motifs are within these windows. **(d)** Chromatin accessibility is most significantly reduced at motif islands containing Zelda and patterning TF motifs. Differential accessibility between *wt* and *zld* embryos was calculated using DESeq2, shown for each island as median log2 fold change values from all time points. Island types that contain more than Zelda motifs show significantly more changes than those with Zelda motifs alone, e.g., the difference between *zld* and *Dl_Zld* islands (p = 8.3e-11, Wilcoxon rank-sum test) and *zld* and *Dl_Twi_Zld* islands (p < 2.22e-16, Wilcoxon rank-sum test).

The results are consistent with Zelda’s role in pioneering, but also reveal the role of patterning TFs. Islands without a Zelda motif typically have very low accessibility and histone modifications, coupled with higher nucleosome occupancy. Islands that only have Zelda motifs and no other motif (Figure 4b, red box) show an increase in chromatin accessibility over time, with an effect proportional to the number of Zelda motifs (Supplemental figure 12d). Although overall, the effect is modest, and these islands have low levels of histone modifications and are not enriched for known developmental enhancers active in blastoderm embryos^73^. By contrast, the highest levels of enhancer accessibility are found at islands that also have motifs for patterning TFs and have the properties of active enhancers. Islands containing motifs for both Zelda and patterning TFs (e.g., Dorsal and Twist) show much higher levels of accessibility, nucleosome depletion, and histone modifications than Zelda-only islands. Interestingly, H3K4me1 correlates better with chromatin accessibility, while H3K27ac correlates better with activity (Figure 4b, Supplemental figure 12c). Taken together, these results suggest that it is the combination of Zelda motifs and patterning TF motifs that generates the highest levels of accessibility, which would explain why it has been challenging to causally link individual TFs such as Bicoid to increased levels of chromatin accessibility beyond those generated by pioneer TFs^56^.

To detect the effect of patterning TFs on chromatin accessibility experimentally, we took advantage of our *zld^-^* ATAC-seq data. Since the patterning TFs require Zelda for binding, any effects that they have on chromatin accessibility should also be lost in *zld^-^* embryos, in addition to the loss of chromatin accessibility caused by Zelda depletion. Thus, we expect that depleting Zelda has a stronger effect on regions with motifs for both Zelda and patterning TFs compared to those with only Zelda motifs. This was indeed the case (Figure 4d). For example, islands with Zelda, Dorsal, and Twist motifs had a much more pronounced fold-change loss in accessibility than Zelda-only islands (p < 2.2e-16, Wilcoxon rank-sum test). These experimental results confirm a model by which high levels of chromatin accessibility are established in a hierarchical manner by a combination of motifs for the pioneer Zelda and the downstream patterning TFs.

### Patterning transcription factors contribute to accessibility when mediating activation

Our results suggest that patterning TFs increase chromatin accessibility when their motifs are present in specific combinations that include Zelda motifs. Enhancers with such motif combinations also tend to be active enhancers, raising the question whether enhancer activity and accessibility are directly functionally coupled. This would be consistent with previous observations that the highest levels of accessibility and TF binding are often found at active enhancers^73–75,77,95,96^. Alternatively, it is possible that the binding of patterning TFs also consistently contributes to the accessibility, but that their dependence on Zelda motifs for binding creates the requirement for motif combinations. The poor correlation between the binding of patterning TFs and their contribution to accessibility argues against this hypothesis (Figure 2f), but we cannot rule out that this is due to limitations of the ChromBPNet model. To distinguish whether patterning TFs mediate increased accessibility through their binding or through their effect on enhancer activity, we leveraged the strengths of *Drosophila* genetics to experimentally test the context-dependent role of Dorsal in chromatin accessibility.

Dorsal is present in the early embryo as a ventral-to-dorsal nuclear concentration gradient that is set up by maternal Toll signaling on the ventral side. At high levels of nuclear Dorsal, the nuclei acquire mesodermal identity; at low levels of Dorsal, they acquire neuroectodermal identity; in the absence of Dorsal, they acquire dorsal ectodermal identity^61^(Figure 5a). The key to Dorsal’s ability to specify three tissue types is its ability to function as a dual transcription factor that can activate mesoderm and neuroectoderm genes and repress dorsal ectoderm genes. This switch in function is possible because the repressed enhancers have Dorsal motifs that are flanked by low-affinity motifs for the repressor Capicua (Cic)^59,97–99^.

**Figure 5.**
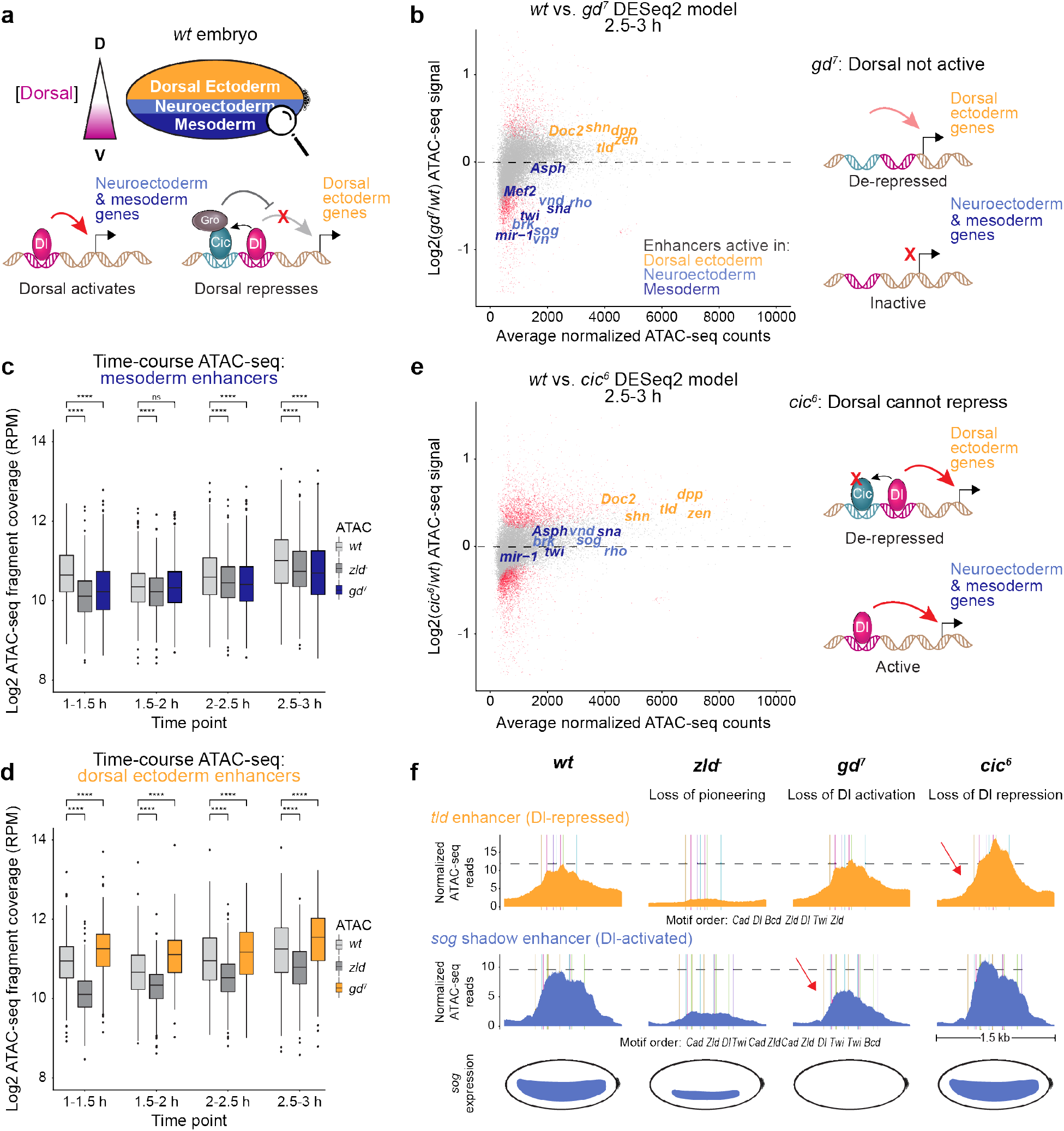
Patterning transcription factors increase chromatin accessibility through transcriptional activation. **(a)** Schematic summary of dorsoventral patterning in the early *Drosophila* embryo. Toll signaling sets up a ventral-to-dorsal nuclear concentration gradient of the maternally-supplied Dorsal TF. High concentrations of Dorsal give rise to mesoderm on the ventral side of the embryo, while low Dorsal concentrations lead to neuroectoderm formation in the lateral regions. Dorsal ectoderm is formed on the dorsal side, where there is no nuclear Dorsal present. In Dorsal-containing tissues (i.e., mesoderm and neuroectoderm), Dorsal is an activator of mesodermal and neuroectodermal target genes but a repressor of dorsal ectodermal genes. Dorsal repression occurs through a cooperative relationship with Capicua, whose low-affinity motifs flank Dorsal motifs in dorsal ectoderm target enhancers. Capicua binding at these regions depends on Dorsal, and it then recruits the corepressor Groucho to repress the dorsal ectoderm genes. **(b)** Chromatin accessibility is specifically reduced at Dorsal-activated enhancers but not at Dorsal-repressed enhancers in embryos lacking nuclear Dorsal. ATAC-seq time course experiments were performed in *gd^7^* embryos, in which Dorsal is not activated and thus represent entirely dorsal ectoderm. Differential accessibility was conducted between *wt* and *gd^7^* embryos for all time points and the MA plot for the 2.5-3 h AEL time point is shown. Red dots represent statistically significant differentially accessible ATAC-seq peaks (FDR = 0.05) and known dorsoventral enhancers are colored by the tissue type in which they are active. Chromatin accessibility is significantly reduced at Dorsal-activated enhancers. Dorsal-repressed enhancers do not lose accessibility in *gd^7^* embryos. **(c)** Mesoderm enhancers lose chromatin accessibility in *gd^7^* embryos. Normalized ATAC-seq fragment coverage from wt, *zld^-^*, and *gd^7^* embryos was calculated at previously determined mesoderm enhancers (n = 416)^102^ across a 1 kb window. Statistical significance was determined between *wt* and *zld^-^* embryos and *wt* and *gd^7^* embryos using Wilcoxon rank-sum tests, where four asterisks is p < 0.0001. In *gd^7^* embryos, mesoderm enhancers are inactive. **(d)** Dorsal ectoderm enhancers gain chromatin accessibility in *gd^7^* embryos. The same analysis used in Figure 5c was performed at dorsal ectoderm enhancers (n = 380). In *gd^7^* embryos, dorsal ectoderm enhancers are active. **(e)** Chromatin accessibility is increased at Dorsal-repressed enhancers upon gaining Dorsal activation. ATAC-seq experiments were performed in *cic^6^* embryos, where Capicua’s interactions with Groucho are abrogated, thus eliminating Dorsal-mediated repression and converting Dorsal into an activator at these enhancers. Differential accessibility analysis between wt and *cic^6^* embryos was performed as in Figure 5b. Dorsal-repressed enhancers, which now gain Dorsal activation, show a significant increase in chromatin accessibility while mesoderm and neuroectoderm enhancers are not differentially accessible. **(f)** Summary of chromatin accessibility at a Dorsal-repressed enhancer (*tld*) and Dorsal-activated enhancer (*sog* shadow) upon loss of Zelda, nuclear Dorsal, and Dorsal-mediated repression. Normalized ATAC-seq fragment coverage is shown from the 2.5-3 h AEL time point across a 1.5 kb window, with the *wt* ATAC-seq maximum value marked with the dotted gray line. Colored bars are BPNet-mapped motifs according to the specified order. The dm6 enhancer coordinates are chr3R: 24,748,748 - 24,750,248 (*tld*) and chrX: 15,646,300 - 15,647,800 (*sog* shadow). Without Zelda, both enhancers dramatically lose accessibility. In *gd^7^* embryos, the Dorsal-activated *sog* shadow enhancer loses chromatin accessibility (red arrow is DESeq2 statistical significance) upon loss of Dorsal activation, while the Dorsal-repressed *tld* enhancer does not lose accessibility (n.s.). In *cic^6^* embryos, the *tld* enhancer gains chromatin accessibility upon gaining Dorsal activation, while the *sog* shadow enhancer shows little change in chromatin accessibility (n.s.). The *sog* expression patterns are based on previous *in situ* hybridization experiments^42,60,104^ and show separate effects from the loss of pioneering and the loss of enhancer activation.

If Dorsal consistently contributes to chromatin accessibility by binding to target enhancers, we would expect that loss of Dorsal leads to decreased chromatin accessibility at all its target genes. To test this, we used *gastrulation defective (gd^7^)* mutant embryos, which are defective in maternal Toll signaling and thus Dorsal remains cytoplasmic and inactive in the entire embryo. As a result, these embryos acquire entirely dorsal ectoderm fate^77,100–102^. After validating the *gd^7^* mutant embryos (Supplemental figure 14a), we performed ATAC-seq time course experiments, producing replicates that were highly correlated (Supplemental figure 13). Using DESeq2^103^, we analyzed the differential accessibility upon loss of Dorsal (*gd^7^*) as compared to wildtype (last time point in Figure 5b, earlier times points in Supplemental figure 14b).

When we examined known Dorsal target enhancers, we noticed a striking difference in accessibility between enhancers that are activated by Dorsal versus those that are repressed. Mesoderm enhancers (e.g., *twi, sna)* and neuroectoderm enhancers (e.g., *sog*, *brk)*, which are activated by Dorsal, show significantly decreased accessibility upon loss of Dorsal (purples in Figure 5b). Conversely, the Dorsal-repressed enhancers do not show decreased accessibility and even show a slight increase, even though they lost Dorsal binding (orange in Figure 5b). These results suggest that Dorsal’s ability to increase chromatin accessibility is tied to its role as transcriptional activator.

To confirm this effect more broadly and over time, we used a set of previously identified enhancers that have differential H3K27ac levels in *gd^7^* mutant embryos and show appropriately regulated target genes nearby^102^. We plotted the ATAC-seq signal for each time point and found that the mesoderm enhancers showed decreased chromatin accessibility in both *zld^-^* and *gd^7^* embryos (Figure 5c). Neuroectodermal enhancers activated by Dorsal show a similar loss in chromatin accessibility (Supplemental figure 14c). Dorsal ectoderm enhancers on the other hand also lose accessibility in *zld^-^* embryos, but instead gain accessibility in *gd^7^* embryos, where they gain activation (Figure 5d). This further corroborates that loss of Dorsal does not always lead to loss of accessibility at Dorsal-bound enhancers, but rather depends on whether Dorsal functions as an activator at these enhancers.

One could argue that loss of Dorsal at dorsal ectoderm enhancers did not lead to a loss of accessibility because other TFs are bound to these regions in *gd^7^* embryos. However, the effect was observed from the earliest time point on, when the primary mechanism of dorsoventral patterning occurs through Dorsal. Enhancers such as *tld*, *zen*, and *dpp* are well studied and known to be regulated by Dorsal repression with the help of Capicua. In *gd^7^* embryos, these enhancers lose both Dorsal and Capicua binding and become de-repressed^59,98,99^. Since we observe a subtle increase in chromatin accessibility, this suggests that chromatin accessibility is tied to enhancer activity, not Dorsal binding.

To test this hypothesis more directly, we specifically manipulated the ability of Dorsal to repress without affecting its ability to activate. In *cic*^6^ mutant embryos, Capicua has a small deletion in its interaction domain (N2) with the co-repressor Groucho and no longer functions as a repressor^59^ (Figure 5e). As a result, Dorsal can still activate mesoderm and neuroectoderm enhancers but it can no longer function as a repressor at dorsal ectodermal enhancers, where it is now expected to function as a weak activator^59^. Thus, in *cic^6^* embryos, the Dorsal-activated enhancers should be unchanged compared to wildtype, while enhancers normally repressed by Dorsal should have higher chromatin accessibility. Indeed, when we performed ATAC-seq experiments in *cic^6^* mutant embryos (Supplemental figure 14d), we found that dorsal ectoderm enhancers showed statistically significant increased accessibility (Figure 5e, orange), while mesoderm and neuroectoderm enhancers not regulated by Capicua generally remained unchanged (Figure 5e, purples). These results demonstrate that the chromatin accessibility at Dorsal target enhancers depends on the activation state induced by Dorsal rather than the binding of Dorsal.

Interestingly, the results also suggest that repressors such as Capicua could decrease chromatin accessibility at their target enhancers. Enhancers that are repressed by Capicua independently of Dorsal through high-affinity Capicua motifs (e.g., *hkb, tll, hb*, and *ind*)^59,105–107^ also increased in accessibility in *cic^6^* mutant embryos, while control enhancers (e.g., *cnc, oc, ems*, and *gt)^59^* remained unchanged (Supplemental figure 14e, f). Whether Capicua directly decreases chromatin accessibility or whether it counteracts the activity of other TFs such as Bicoid and Caudal remains to be tested.

In summary, our results suggest that chromatin accessibility levels depend on both pioneering and enhancer activation. Pioneering by Zelda consistently contributes to accessibility, while the effect of patterning TFs such as Dorsal is context-dependent. This is well illustrated at the Dorsal-repressed enhancer *tld*^59^ and the Dorsal-activated *sog* shadow enhancer^42^ (Figure 5f). In both cases, the chromatin accessibility is dramatically reduced in *zld^-^* embryos due to the loss of pioneering (Figure 5f, second panel). Loss of Dorsal (*gd^7^*) led to a modest but significant decrease in accessibility across the Dorsal-activated enhancer, while the Dorsal-repressed enhancer showed little change (Figure 5f, third panel). Converting Dorsal from a repressor into an activator (*cic^6^*), caused a significant increase in chromatin accessibility across the Dorsal-repressed enhancer, while accessibility was essentially unchanged across the Dorsal-activated enhancer (Figure 5f, fourth panel). The same patterns were observed at other enhancers, including those for *dpp* and *sna* (Supplemental figure 14g), confirming the distinct roles of Zelda and Dorsal.

Pioneering and enhancer activation do not simply differ because of different effect sizes, but rather appear to be distinct processes. While chromatin accessibility is more dramatically affected by the loss of Zelda than the loss of Dorsal, the inverse is true for the effect on gene expression. In the absence of Dorsal, the expression of *sog* is completely abolished^104^, while in the absence of Zelda, *sog* expression is delayed and narrowed but still occurs with high concentrations of Dorsal^42,60^. Thus, Zelda has a stronger effect on chromatin accessibility, while Dorsal has a stronger effect on activation, arguing that they involve functionally separable processes that both have effects on chromatin accessibility.

## Discussion

Here, through combining TF binding data, chromatin accessibility data, deep learning models capable of learning both datasets independently of one another, and using classic *Drosophila* genetics as a validation tool, we asked how TFs mediate chromatin accessibility in the *Drosophila* embryo. We investigated whether the role of opening chromatin is restricted to TFs axiomatically classified as pioneers or if TFs more generally contribute to chromatin accessibility. We uncovered the cis-regulatory sequence rules and distinct underlying mechanisms of this process.

Our results suggest a hierarchical two-tier model, where chromatin accessibility is established first through pioneering but is further increased during enhancer activation (Figure 6). Importantly, the sequence rules for chromatin accessibility during activation are distinct from those that mediate pioneering. Pioneers like Zelda are the first to bind to their motifs genome-wide and consistently bestow basal accessibility by reading out motif affinity, thereby creating a more permissive landscape for other TFs. In contrast, the patterning TFs require an already accessible state for their binding and increase chromatin accessibility in a contextdependent manner since they only increase chromatin accessibility when mediating enhancer activation. For example, when Dorsal motifs are flanked by motifs for the repressor Capicua in dorsal ectoderm enhancers, no increase in chromatin accessibility is observed. These enhancers do however show an increase in chromatin accessibility when Capicua is mutated such that Dorsal can no longer repress and instead becomes an activator. This demonstrates that the increase in accessibility is not dependent on Dorsal binding *per se* but on the total effect that the TFs have on the activation of the enhancer, and thus is governed by the cis-regulatory rules of activation. This contrasts with Zelda, which consistently increases chromatin accessibility in the absence of enhancer activation.

**Figure 6.**
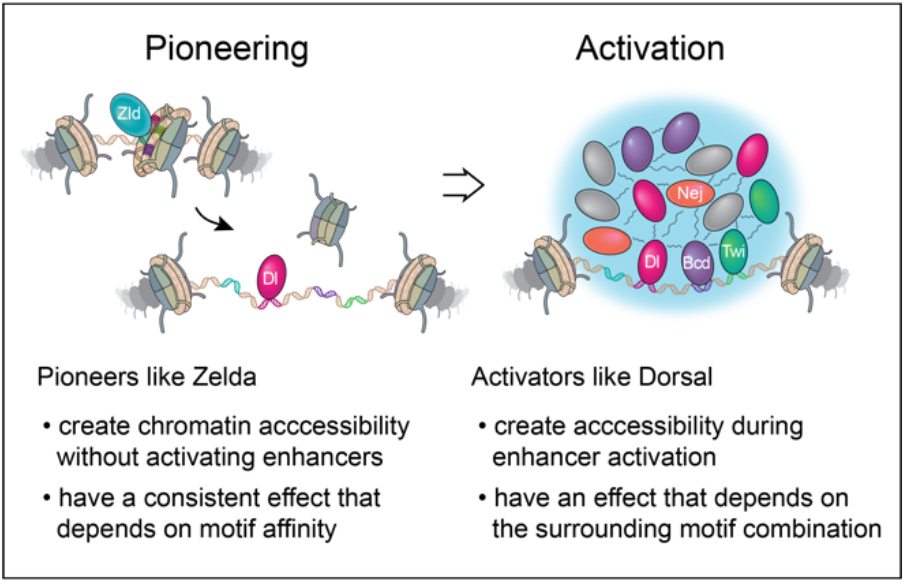
Pioneering and enhancer activation increase chromatin accessibility. Chromatin accessibility at enhancers is established in a two-tier process that involves pioneering and activation. First, the pioneer Zelda bestows basal chromatin accessibility at enhancers, without activating them, by reading out its motif affinity on nucleosomal DNA. Zelda’s pioneering is a consistent effect that is not dependent on the combination of motifs in the enhancer. The pioneering then allows the binding of patterning TFs such as Dorsal, which require an accessible state of the DNA to bind to their motifs. Hubs may then form on accessible regions when sufficient concentrations of patterning TFs bind and interact with each other and co-factors through multivalent weak interactions. In this way, hub formation is DNA-templated and facilitated by Zelda’s global pioneering. Whether or not Zelda is also present in these hubs is unclear. During enhancer activation, chromatin accessibility is further increased, perhaps by hubs recruiting Nej, the *Drosophila* CBP, which mediates histone acetylation at enhancers. Since the TF hubs appear dynamic, they could leave the DNA and make the region more accessible.

The functional separation between pioneering and activation is consistent with previous observations in the early *Drosophila* embryo. Zelda unambiguously generates chromatin accessibility very early on, but is insufficient for the activation of most enhancers and functions together with patterning TFs during zygotic genome activation^40,76,108–110^. At many enhancers, Zelda is not even strictly required for enhancer activation since many patterning genes eventually become expressed in *zld^-^* embryos^38^. Zelda is however a strong potentiator of transcription^5,42,44,45,60^. This suggests that Zelda’s effect on chromatin accessibility is not required for activation but boosts the effect of activators. A similar potentiating effect of Zelda has been observed at the level of transcriptional bursting. Dorsal mainly affected the burst frequency, while Zelda had an additional effect on the burst size^108^.

These functional differences are consistent with pioneering and activation being physically separate processes. Zelda binds its motifs in the presence of nucleosomes^19,48^, while Dorsal, Twist, Caudal, and Bicoid require accessible DNA for binding^5,6,42,44,45,47,60^. Consistent with Zelda binding nucleosomes *in vivo*, Zelda has a broad binding footprint in ChIP-nexus data (Figure 1c), which could be mediated by indirect contacts to DNA through nucleosomes. In contrast, patterning TFs have sharper and narrower footprints consistent with their binding primarily accessible genomic DNA (Figure 1c). While Zelda could also bind to accessible regions, this may not occur to a large extent since Zelda binds to chromatin in a rapid and transient manner^46^ and does not co-localize with Pol II or at sites of active transcription^46,109^. Thus, pioneering appears to be the process associated with nucleosome removal, while enhancer activation occurs on accessible DNA.

How could motifs mediate pioneering? Studies *in vivo* all point to a constant involvement of ATP-dependent chromatin remodeling^111,112^, but how pioneer TFs recognize their motifs on nucleosomal DNA and interact with chromatin remodelers is not clear. TF binding to nucleosomes *in vitro* tends to be structurally restricted and can be preferred at certain positions on the nucleosome^19,20,82,83^, but it is unclear whether these structural restrictions are relevant *in vivo*. Our finding that Zelda very precisely reads out motif affinity and commensurately increases chromatin accessibility is therefore remarkable. While we cannot rule out that the influence of nucleosome position was not learned by ChromBPNet, our results argue against a strong dependence on the motif’s relative position on the nucleosome. This is also consistent with our previous study, where we did not find a preferred position for Zelda motifs on *in vivo* nucleosomes^6^. Instead, our results argue that pioneer TFs recognize their motifs *in vivo* more efficiently than *in vitro*, perhaps aided by chromatin remodelers.

How could enhancer activation occur dependent on sequence contexts? Since enhancer activation depends on the motif combination and accessible DNA, we propose that it occurs through DNA-mediated hub formation (Figure 6). When DNA with a set of motifs become accessible and bound by TFs and co-factors, the DNA serves as a seed to induce surface condensation, which locally concentrates the proteins into hubs^113–115^. In support of this model, hubs have been observed via imaging studies for multiple TFs in the early *Drosophila* embryo, including Zelda, Dorsal, and Bicoid^46,47,60,109^. Hubs containing either Dorsal or Bicoid were dependent on Zelda, which is consistent with DNA accessibility being a requisite for hub formation. Furthermore, Dorsal and Bicoid have been reported to recruit the co-factor Nej, the *Drosophila* CBP^104,116–118^, which could promote hub formation. Lastly, if hubs regulate transcriptional bursting, this could explain why Dorsal and Zelda have different effects. Dorsal may determine the burst frequency by regulating the speed of hub formation on already accessible DNA, while Zelda also facilitates chromatin accessibility and thus may affect the burst size by providing more time and space for hub formation. While this hub model fits well with current data, it does not explain how activation increases the accessibility further. Further studies are needed to better understand the role of hubs.

Our results suggest that the relationship between accessible DNA, TF binding, and enhancer activation is more complex than previously thought. Notably, we found that our deep learning models correctly identified the motif for the pioneer TF GAF to play a strong role in chromatin accessibility, but our models also predicted that GAF does not play the same role as Zelda in helping other TFs bind. While GAF is predicted to boost its own binding, it does not seem to strongly promote the binding of the patterning TFs. One explanation for the difference may be the residence time on DNA. While Zelda binds DNA only transiently on the order of seconds^46^, GAF multimerizes on DNA and remains on chromatin on the order of minutes^119–122^. Such stable binding makes sense in the light of GAF’s role in 3D genome structure^122–126^ and transcriptional memory^120,127,128^. Thus, GAF could generate accessible chromatin but by binding to the newly opened DNA itself, it could partially occlude the binding of additional TFs. These results suggest that accessibility does not necessarily make the region accessible for all other TFs and further highlight that accessibility is not always a perfect proxy for activation.

A separate contribution of pioneering and enhancer activation towards chromatin accessibility likely applies to mammals. In mammals, the highest accessibility is typically also found at active enhancers^129–132^, yet chromatin accessibility is often only a mediocre predictor for enhancer activity^133–135^. Without TF binding data and prior knowledge, it can however be difficult to deduce from accessibility data alone whether a TF bestows chromatin accessibility as a pioneer TF, as an activator, or both^16,17,136^. For example, in our later time points where Dorsal binding is highest, Dorsal more consistently promotes chromatin accessibility (Figure 2f), thus behaving more like a pioneer. It might therefore initially require a combined approach, which includes TF binding, chromatin accessibility, deep learning, and additional experiments, to better distinguish between the mammalian TFs that drive chromatin accessibility and those that follow it.

## Supplemental files

Supplemental file 1: BPNet-identified and mapped motifs for Zelda, GAF, Bicoid, Caudal, Dorsal, and Twist. Motif coordinates come from the *Drosophila melanogaster* dm6 genome assembly.

Supplemental file 2: Motif islands, with provided coordinates aligned to the *Drosophila melanogaster* dm6 genome assembly. Islands were tested for overlaps with known active enhancers^73^. The normalized ATAC-seq signal calculated 250 bp across each island is provided from wildtype, *gd^7^, zld^-^*, and *cic^6^* embryos. Island types with fewer than 30 instances were excluded.

## Data and code availability

The raw and processed data for ChIP-nexus, ChIP-seq, ATAC-seq, MNase-seq and protein binding microarray experiments are available from GEO under series accession number GSE218852. All code used to process and analyze the data can be accessed at https://github.com/zeitlingerlab/Brennan_Zelda_2023. The ChIP-nexus protocol and the data processing description can be found at https://research.stowers.org/zeitlingerlab/protocols.html.

Trained BPNet and ChromBPNet models will be available at Zenodo and Kipoi following review. Original data, including microscopy images, can be accessed from the Stowers Original Data Repository at http://www.stowers.org/research/publications/libpb-2357.

## Acknowledgements

We thank Žiga Avsec, Robb Krumlauf, Kausik Si, Vikki Weake, and Zeitlinger lab members for helpful comments and suggestions on the manuscript. We thank Mark Miller for help with figure illustrations, Beth Canfield for help with *Drosophila* husbandry, and the following Stowers Institute core facilities for their support: Sequencing and Discovery Genomics (Anoja Perera, Michael Peterson, and Amanda Lawlor), Lab Services (Stacey Walker), Cytometry (Jeff Haug and KyeongMin Bae), and Computational Biology (Hua Li, Madelaine Gogol, and Jonathon Russell). This research reported in this publication was supported by the Stowers Institute for Medical research and by the Eunice Kennedy Shriver National Institute of Child Health & Human Development of the National Institutes of Health under the F31 Award Number F31HD108901 to K.J.B. The content is solely the responsibility of the authors and does not necessarily represent the official views of the National Institutes of Health.

## Author contributions

K.J.B. and J.Z. conceived the project as part of K.J.B.’s thesis research to fulfill the requirements for the Graduate School of the Stowers Institute for Medical Research. K.J.B. and J.Z. designed the genomics and genetics experiments, which were performed by K.J.B. and S.K. Protein binding microarray experiments were designed by H-Y.L., A.W.H.Y., T.R.H., and C.A.R. and performed by H-Y.L., and A.W.H.Y. Computational methods were conceived and designed by M.W., A.P., A.K., and J.Z. Deep learning model training, computational analysis, and *in silico* experiments were performed by M.W. Additional genomics data analyses were done by M.W. and K.J.B. The manuscript was prepared by K.J.B., M.W., and J.Z. with input from all authors.

## Methods

### *Drosophila* strains

Oregon-R flies were used as the wildtype *(wt)* strain in all experiments. Embryos depleted for maternal Zelda *(zld^-^*) were generated by crossing *UAS-shRNA-zld* females to *MTD-Gal4* males as previously described^6^ and tested for embryonic lethality^38^ and Zelda depletion using immunostaining (Figure 3). Embryos lacking nuclear Dorsal were laid by *gd^7^/gd^7^* mothers generated from a *gd^7^/winscy, P{hs-hid}5* stock that was heat-shocked at the larval stage at 37°C for 1 hour on two consecutive days to eliminate heterozygous mothers^6^. Loss of the hs-hid sequence was confirmed using PCR on genomic DNA extracted from heat-shock survivors. The *cic^6^/TM3, Sb^1^* stock was generated using CRISPR/Cas9 as previously described^59^. *Cic6* embryos were collected from *cic^6^/cic^6^* mothers identified by *wt* bristles and were confirmed to be embryonic lethal.

### Embryo collections, fixation, and sorting

All embryos were collected from population cages using apple juice plates with yeast paste, following two pre-clearings as previously described^58,137^. For ChIP-nexus, ChIP-seq, and MNase-seq experiments, embryos were collected for 1 h and aged for 2 h at 25°C, yielding collections of 2-3 h after egg laying (AEL). For ATAC-seq, embryos were collected in 30-minute windows and aged accordingly to generate the 1-1.5, 1.5-2, 2-2.5, and 2.5-3 h AEL time points. All embryos were dechorionated using 50% bleach for 2 minutes and sufficiently rinsed with water afterwards. For ATAC-seq, embryos were hand-sorted based on morphology in ice-cold PBT immediately following dechorionation using an inverted contrasting microscope (Leica DMIL) as described^137^. For ChIP-nexus, ChIP-seq, and MNase-seq, embryos were first fixed with 1.8% formaldehyde in heptane and embryo fix buffer (50 mM HEPES, 1 mM EDTA, 0.5 mM EGTA, 100 mM NaCl) while vortexing for 15 minutes. For ChIP-nexus and ChIP-seq, the vitelline membrane was removed using methanol/heptane and embryos were stored in methanol at −20°C until use. For these experiments, embryos were rehydrated using PBT and sorted to remove out-of-stage embryos using either hand-sorting or cytometry (Copas Plus, macroparticle sorter, Union Biometrica). For MNase-seq, embryos were spun down at 500 x g, 4°C, for 1 minute, and fixation was quenched by adding 10 mL PBT-glycine (125 mM glycine in PBT) and vortexing for 2 minutes. Embryos were hand-sorted based on morphology in ice-cold PBT and then used in MNase-seq experiments.

### ChIP-nexus and ChIP-seq experiments

For each ChIP, 10 μg of antibody was coupled to 50 μL of Protein A Dynabeads (Invitrogen) and incubated overnight at 4°C prior to ChIP. All ChIP-nexus experiments were performed using antibodies custom generated by Genscript: Zelda (aa 1117-1327), Dorsal (aa 39-346), Twist (C-terminus), Bicoid (C-terminus), Caudal (aa 1-214), GAF (aa 13-82). ChIP-seq experiments were performed with the following commercially available antibodies: H3K27ac (Active motif, 39133) and H3K4me1 (Active motif, 39635). For all TFs, at least three biological replicates were performed using embryos from different collections. For ChIP-seq, at least two biological replicates were performed in the same way. Approximately 0.2-0.4 grams of fixed 2-3 h AEL embryos were used for all ChIP experiments. Chromatin extracts were prepared by douncing embryos in Lysis Buffer A1 (15 mM HEPES pH 7.5, 15 mM NaCl, 60 mM KCl, 4 mM MgCl2, 0.5% Triton X-100, 0.5 mM DTT (add fresh)), washing nuclei with ChIP Buffer A2 (15 mM HEPES pH 7.5, 140 mM NaCl, 1 mM EDTA, 0.5 mM EGTA, 1% Triton X-100, 0.5% N-lauroylsarcosine, 0.1% sodium deoxycholate, and 0.1% SDS), and sonicating with a Bioruptor Pico (Diagenode) for six cycles of 30 seconds on and 30 seconds off. ChIP-nexus library preparation was performed as previously described^58^, except that the ChIP-nexus adapter mix contained four fixed barcodes and PCR library amplification was performed directly after circularization of the purified DNA fragments (without addition of the oligo and BamHI digestion). ChIP-seq was performed as previously described and included a whole cell extract (WCE)^68,77^. Single-end sequencing was performed on an Illumina NextSeq 500 instrument (75 or 150 cycles). The full ChIP-nexus protocol can be found on the Zeitlinger lab website at https://research.stowers.org/zeitlingerlab/protocols.html.

### ATAC-seq experiments

For ATAC-seq time course experiments, the following amounts of hand-sorted embryos were used: 400 embryos (1-1.5 h AEL); 100 embryos (1.5-2 h AEL); 40 embryos (2-2.5 h AEL, 2.5-3 h AEL). Following sorting, embryos were immediately dounced in ATAC Resuspension Buffer (10 mM Tris-HCl pH 7.4, 10 mM NaCl, 3 mM MgCl2) with 0.1% IGEPAL CA-630 and nuclei were harvested by centrifugation. Tn5 transposition was performed as previously described^71,72^. Briefly, the nuclear pellet was incubated for 3 minutes on ice in ATAC resuspension buffer supplemented with 0.1% IGEPAL CA-630, 0.1% Tween-20, and 0.01% Digitonin (Promega, G9441). The reaction was stopped by adding ATAC Resuspension Buffer with 0.1% Tween-20 followed by centrifugation. Tagmentation took place at 37°C for 30 minutes at 1000 rpm in a 50 μL reaction volume containing 10 μL of 5x Tagment DNA Buffer (50 mM Tris-HCl pH 7.4, 25 mM MgCl_2_, 50% DMF) 16.5 μL 1x PBS, 0.5 μL 10% Tween-20, 0.5 μL 1% Digitonin,1-2 μM assembled transposome and water. Tn5 transposome was purified in-house as previously described^138^. The resulting fragments were purified using the Monarch PCR & DNA Cleanup Kit (NEB). Libraries were constructed using Illumina Nextera Dual Indexing, and qPCR was used to prevent over-amplification as described^72^. At least three biological replicates were generated and paired-end sequencing was performed on an Illumina NextSeq 500 instrument (2x 75 bp cycles).

### MNase-seq experiments

For each MNase digestion, 100 hand-sorted 2-3 h AEL Drosophila embryos were used. Nuclei were extracted by douncing in PBS with 0.1% IGEPAL CA-630. The nuclei were harvested by centrifugation and resuspended gently in MNase Digestion Buffer (PBS with 0.1% Triton X-100 and 1 mM CaCl_2_). MNase digestion was performed with 100 U MNase (NEB, M0247S) for 30 minutes at 37°C. The reaction was stopped with 20 mM EGTA. The nuclei were treated with 50 μg/ml RNase A (Thermo Scientific, EN0531) for 1 hour at 37 °C and 1000 rpm, and subsequently incubated overnight at 65 °C and 1000 rpm with 200 μg/ml Proteinase K (Invitrogen, 100005393) and 0.5% SDS for reverse crosslinking. DNA was extracted using phenol-chloroform (VWR, K169). Libraries were constructed from 10 ng purified DNA using the High Throughput Library Prep Kit from KAPA Biosystems (KK8234) according to the manufacturer’s instructions. Three experimental replicates were performed. Paired-end sequencing was performed on an Illumina NextSeq 500 instrument (2x 75 bp cycles).

### Antibody staining and microscopy experiments

Embryos were collected and aged to be 2-3 hours old, fixed with 1.8% formaldehyde, and stored in 100% methanol at −20°C prior to immunostaining. Embryo aliquots were rehydrated in an ethanol:PBT gradient and blocked for 30 minutes using the Roche Western blocking reagent (11921681001) and PBT. Primary antibody incubation occurred at 4°C overnight with a 1:200 antibody dilution in PBT/blocking reagent with the same Zelda, Dorsal, and Twist antibodies used for ChIP-nexus experiments. Embryos were then washed six times with PBT, blocked again, and incubated with a donkey anti-rabbit IgG Alexa Fluor 568 secondary antibody (Thermo Fisher, A10042), 1:500, at 4°C overnight. After eight washes with PBT, embryos were mounted with ProLong Gold Antifade Mountant with DAPI (Invitrogen, P36931). Images were acquired on a Zeiss LSM-780 laser scanning confocal microscope with a 32 channel GaAsP detector and a plan-apochromat 10x objective lens, N.A. 0.45, using the ZEN Black 2.3 SP1 software by Zeiss. The Alexa Fluor 568 track used a DPSS 561 nm laser excitation at 6.5%, and the DAPI track used a Diode 405 nm laser excitation at 6.0%. Images were collected using a frame size of 1024 x 1024, a zoom of 1.5, and a pixel dwell time of 3.15 μs. Confocal z-stacks were maximum intensity projected and all image processing steps were performed using FIJI^139^. All microscopy and processing settings were kept the same when comparing *wt* to *zld^-^* or *gd^7^* embryos.

### Protein binding microarray experiments

For all PBM experiments, the C-terminal region of Zelda, which includes the four zinc fingers (#3-6) that are known to bind CAGGTAG motifs, were used^37,38^. These zinc fingers were cloned into a T7-driven GST expression vector, pTH6838. The TF sample was expressed by using a PURExpress *In Vitro* Protein Synthesis Kit (New England BioLabs) and analyzed in duplicate on two different PBM arrays (HK and ME) with differing probe sequences. PBM laboratory methods including data analysis followed the procedures previously described^140,141^. PBM data were generated with motifs derived using Top10AlignZ^88^. Z-scores and E-scores were calculated for each 8-mer as previously described^87,88^. Octamers were grouped together based on their heptad sequences while also considering reverse complements, and the median E-score and Z-score was calculated for each 7-mer. The heptad sequences matching BPNet-mapped Zelda motifs were then extracted and the two PBM replicates were averaged for each Zelda motif.

### ChIP-nexus data processing

ChIP-nexus single-end sequencing reads were preprocessed by trimming off fixed and random barcodes and reassigning them to FASTQ read names. ChIP-nexus adapter fragments were trimmed from the 3’ end of the fragments using cutadapt (v.2.5^142^). ChIP-nexus reads were aligned using bowtie2 (v.2.3.5.1^143^) to the *Drosophila melanogaster* genome assembly dm6. Aligned ChIP-nexus BAM files were deduplicated based on unique fragment coordinates and barcode assignments. Normalized ChIP-nexus coverage was acquired through reads-per-million (RPM) normalization, where the ChIP-nexus sample coverage was scaled by the total number of reads divided by 10^6^. ChIP-nexus peaks were mapped using MACS2 (v.2.2.7.1^144^) with parameters designed to resimulate the full fragment length coverage rather than the single stop base coverage (--keep-dup=all - f=BAM --shift=-75 --extsize=150). ChIP-nexus peaks were filtered for pairwise reproducibility using the Irreproducible Discovery Rate framework (IDR) (v.2.0.3^145^). Peaks used for downstream analysis were selected from the largest pairwise comparison using the IDR framework.

### ATAC-seq data processing

ATAC-seq paired-end sequencing reads were aligned using bowtie2 (v.2.3.5.1^143^) to the *Drosophila melanogaster* genome assembly dm6. Aligned ATAC-seq BAM files were marked for duplicates using Picard (v.2.23.8^146^) based on unique fragment coordinates, deduplicated, reoriented according to a Tn5 enzymatic cut correction of −4/+4 on fragment ends, filtered to contain fragment lengths no greater than 600 bp, and corrected for dovetailed reads. Normalized ATAC-seq coverage was acquired through reads-per-million (RPM) normalization, where the ATAC-seq sample coverage was scaled by the total number of reads divided by 10^6^, as performed previously^50,76^. Cut site ATAC-seq coverage was acquired by treating each of the fragment ends as a “cut event” and generating coverage based on only these “cut events”. ATAC-seq peaks were mapped using MACS2 (v.2.2.7.1^144^) with default paired-end parameters using ATAC-seq fragment coverage. ATAC-seq peaks were filtered for pairwise reproducibility using the Irreproducible Discovery Rate framework (IDR) (v.2.0.3^145^). Peaks used for downstream analysis were selected from the largest pairwise comparison using the IDR framework.

### ChIP-seq data processing

ChIP-seq single-end sequencing reads were aligned using bowtie2 (v.2.3.5.1^143^) to the *Drosophila melanogaster* genome assembly dm6. Aligned ChIP-seq BAM files were deduplicated based on unique fragment coordinates and fragments extended based on the average experiment fragment length as determined with an Agilent 2100 Bioanalyzer. Normalized ChIP-seq coverage was acquired using the deepTools subfeature bamCompare (v.3.5.1^147^) using parameters to generate log2 fold-change scaling (--scaleFactorsMethod=readCount --operation=log2 --binSize=1). ChIP-seq peaks were mapped using MACS2 (v.2.2.7.1^144^) with default parameters and an applied background coverage using the associated WCE ChIP-seq control experiment. ChIP-seq peaks were filtered for pairwise reproducibility using the Irreproducible Discovery Rate framework (IDR) (v.2.0.3^145^).

### MNase-seq data processing

MNase-seq paired-end sequencing reads were aligned using bowtie2 (v.2.3.5.1^143^) to the *Drosophila melanogaster* genome assembly dm6. Aligned MNase-seq BAM files were deduplicated based on unique fragment coordinates and filtered to contain fragment lengths no greater than 600 bp. Normalized MNase-seq coverage was acquired through reads-per-million (RPM) normalization, where the MNase-seq sample coverage was scaled by the total number of reads divided by 10^6^.

### BPNet model training and optimization

BPNet architecture and software was applied as previously described^57^. Model inputs were 1000 bp genomic sequences centered on the ChIP-nexus peaks of TFs of interest. Model outputs were the predicted counts (total reads across each region) and predicted profile (coverage signal across each region) for Zelda, Dorsal, Twist, Caudal, Bicoid, and GAF ChIP-nexus experiments. 95,282 IDR-reproducible peaks from Zelda, Dorsal, Twist, Caudal, Bicoid, and GAF ChIP-nexus experiments were pooled and used as model inputs. Validation datasets were peaks located across chr2L (~18% of peaks), test datasets were peaks located across chrX (~19% of peaks), and peaks located across chrY and nonstandard chromosome contigs were excluded from analysis. The remaining regions were used for model training. Hyper-parameters were optimized by selected testing of parameter values deviating from the default BPNet architecture (number of dilational convolutional layers, number of filters in each convolutional layer, filter length of the first convolutional layer, filter length of the deconvolutional layer, learning rate, and counts-to-profile loss balancing). Model optimality was assessed based on counts and profile performance of each task, with a focused emphasis on the Zelda task performance, as this was our key TF of interest. After optimization, the final BPNet model architecture contained 9 dilated convolutional layers, 256 filters in each convolutional layer, a filter length of 7bp for both the input convolutional layer and output deconvolutional layer, a learning rate of 0.004, and a counts-to-profile weighting value (lambda) of 100. Final optimized model performance was assessed through comparing (1) area under the Precision-Recall Curves (auPRC) for profiles over different bins of resolution between observed ChIP-nexus profiles and predicted BPNet profiles (Supplemental figure 2a) and (2) counts correlations of observed ChIP-nexus signals to predicted BPNet signals for each TF (Supplemental figure 2b) as previously described^57^. The auPRC values were benchmarked alongside replicate-replicate, observed-random, and observed-average observed profile comparisons to establish an in-context understanding of predicted profile accuracy. In order to test the stability of this optimized model architecture (fold 1), we trained two additional models with shuffled training, validation, and test sets (three-fold validation). The stability of the performance metrics as well as the stability of the returned downstream motif grammar was compared to the original optimized model training event (Supplemental figure 2c). All BPNet models were implemented and trained using Keras (v2.2.4^148^), TensorFlow1 backend (v.1.7^149^), the Adam optimizer^150^. Training was performed using a NVIDIA^®^ TITAN RTX GPU with CUDA v9.0 and cuDNN v7.0.5 drivers.

### Motif extraction, motif curation, and motif island generation

DeepLIFT (v0.6.9.0, derived from the Kundaje Lab fork of DeepExplain (https://github.com/kundajelab/DeepExplain)^151^ was applied to the trained BPNet model to generate the contribution of each base across a given input sequence to the predicted output counts and profile signals. Contribution scores for counts and profile outputs were generated for all 6 TF tasks. TF-MoDISco (v.0.5.3.0^152^) was then applied across each TF separately. For each TF, regions of high counts contribution were identified, clustered based on within-group contribution and sequence similarity, and consolidated into motifs. The Zelda, Dorsal, Twist, Caudal, Bicoid, and GAF motifs were manually identified based on similarity to previous literature and validation of ChIP-nexus binding from the pertinent TF. Once motifs were characterized and confirmed, they were remapped back to their TF-specific peaks based on both Jaccardian similarity to the TF-MoDISco contribution weight matrix (CWM) and sufficient total absolute contribution across the mapped motif. This mapping approach is previously described^57^. However, as we were interested in lower affinity motif representations than were previously identified by BPNet, mapping thresholds were lowered to mapping the motif if the CWM Jaccard similarity percentile was equal to or greater than 10% and if the total absolute contribution percentile was equal to or greater than 0.5%. After mapping, motifs were filtered for redundant assignment of palindromic sequences and overlapping peaks. Mapped and bound motifs were next clustered into ‘motif islands’ based on their proximity. Each island initially starts as a 200 bp region centered on the motif and gets clustered and merged with another nearby motif island if they overlap. In this manner, islands get extended as long as there is a motif within less than 200 bp. In the end, the vast majority of islands are still between 200-400 bp in width (Supplemental figure 12b). Island types with fewer than 30 genomic instances were filtered out (Supplemental figure 12a). These island clusters were then grouped for downstream analysis.

### ChromBPNet model training and optimization

ChromBPNet is a modification of BPNet, designed to explain the relationship between genomic sequence and base-resolution ATAC-seq cut site coverage^28,79^. ChromBPNet possesses similar model architecture to BPNet, but the training process contains extra steps to accommodate for the Tn5 sequence bias that influences the positions of the ATAC-seq cut sites. If the Tn5 sequence is not accounted for, the positional information of the cut sites cannot be reliably interpreted. ChromBPNet handles this during the training step by simultaneously passing sequence information through (1) a frozen, pre-trained model that has already learned Tn5 sequence bias and (2) an unfrozen, randomly-initialized model that will learn the unbiased sequence rules associated with ATAC-seq cut site coverage. During training, the sequence information will pass through both of these models and their respective outputs will be added together to represent training loss. By adding the two model outputs, ChromBPNet is evaluating both Tn5 sequence bias and sequence rules of accessibility, which can be compared to the actual ATAC-seq cut site coverage (which also possesses both of these features). After the training step has been completed, we remove the frozen Tn5 bias model and apply downstream interpretations only to the second model which contains the unbiased sequence rules that explain accessibility coverage of ATAC-seq cut sites.

To train the highest-quality set of models in the *Drosophila* genome, we trained a custom Tn5 bias model to represent the Tn5 sequence bias in our data. The Tn5 bias model architecture followed ChromBPNet defaults^79^. The Tn5 bias model output was the pooled coverage of the 2.5-3 h ATAC-seq experiments. This time point was chosen for the bias model because it was the most likely time in which this model could have learned underlying sequence grammar of interest and therefore the most optimal to validate against. The Tn5 bias model inputs were genomic regions that met the following critters: (1) closed (non-peak ATAC-seq regions across all time points), (2) unbound (non-peak ChIP-nexus regions across all TFs described above), (3) low-coverage regions (containing less than five times the cut sites as the lowest coverage 2.5-3 h ATAC-seq IDR-reproducible peak region), (4) 2114 bp in width, and (5) at least 750bp away from an annotated fly TSS. These criteria were applied in order to ensure that Tn5 sequence bias was only learned at regions that were closed, inactive, and representative of noise-based cut site coverage. After application of these criteria, the Tn5 bias model was trained on 2,326 training regions and 883 validation regions. Training, validation, and test regions were determined based on the chromosomes reported above for BPNet. In order to validate that the Tn5 bias model learned only Tn5 sequence bias and no other grammar rules, particularly motif-driven rules, we collected Tn5 counts and Tn5 profile contribution scores using the DeepSHAP implementation of DeepLIFT (https://github.com/kundajelab/shap)^151^ and ran TF-MoDISco (v.0.5.16.0^152^). For profile contribution, the Tn5 sequence bias was returned as multiple different logos (Supplemental figure 6b), but no motif consensus logos were returned. For counts contribution, neither Tn5 nor motif consensus logos were returned. This confirmed that our Tn5 bias model was only learning positional Tn5 sequence bias information. In order to follow-up this validation, we injected the sequences of likely canonical motifs into 256 genomic sequences from the test chromosome (chrX) and averaged the effects to observe that the Tn5 bias model did not predict an increase in coverage magnitude (Supplemental figure 6a).

After Tn5 bias model training, ChromBPNet architecture and software (https://github.com/kundajelab/chrombpnet) was applied as described^79^. Model inputs were 2114 bp genomic sequences centered on IDR-reproducible ATAC-seq peaks. In order to fairly compare the results between four ChromBPNet models for each developmental time point measured using ATAC-seq (1-1.5 h, 1.5-2 h, 2-2.5 h, 2.5-3 h), we sought to train each of the models with the pooled IDR-reproducible ATAC-seq peaks from every time point measured. Additionally, because we wished to characterize enhancer accessibility rules, we removed peaks that were within 750 bp of an annotated TSS, as we know that accessibility at promoters can be dictated by different sequence rules than at enhancers. After the time points were pooled and promoter-proximal peaks removed, 41,497 ATAC-seq peaks were included. In order to train more robust models, we also included curated non-peak regions (described above) sampled to 10% of the ATAC-seq peaks for training (4,150 non-peak regions). The inclusion of both peak and non-peak ATAC-seq regions allows the model to better differentiate between accessible and inaccessible sequences. In total, 45,647 regions were used as ChromBPNet model inputs. Validation datasets were peaks located across chr2L (~16% of peaks), test datasets were peaks located across chrX (~19% of peaks), and peaks located across chrY and nonstandard chromosome contigs were excluded from analysis. The remaining regions were used for model training. In addition to shared peaks across different ChromBPNet models to maintain inter-model stability, we also sought to train each of the models with the same ChromBPNet architecture. For this, an optimization search was required, and we again decided to optimize on the pooled coverage of the 2.5-3 h ATAC-seq experiments through selected testing of parameter values deviating from the default ChromBPNet architecture (number of filters in each convolutional layer, filter length of the first convolutional layer, and filter length of the deconvolutional layer). Model optimality was assessed based on the counts and profile performance of the bias-removed predictions, as well as prioritizing model depth to avoid over-distribution of motif grammar within sequence representations. After optimization, the final ChromBPNet model architecture contained 128 filters in each convolutional layer and a filter length of 7 bp for both the input convolutional layer and 75 bp for the output deconvolutional layer. We then trained ChromBPNet models on the pooled cut site coverage of the four developmental time point ATAC-seq experiments (1-1.5 h, 1.5-2 h, 2-2.5 h, 2.5-3 h). Final optimized model performance was assessed through comparing (1) the ability of the model to differentiate peak and non-peak regions using area under the receiver operating characteristic curve (ROC AUC) (Supplemental figure 6c), (2) counts correlations of observed ATAC-seq cut sites to ChromBPNet predictions (Supplemental figure 6d), and (3) profile prediction accuracy of observed ATAC-seq cut sites to ChromBPNet predictions using Jensen-Shannon distances benchmarked by randomly shuffled region profiles (Supplemental figure 6e). In order to test the stability of these different ChromBPNet models, we trained two additional models across each ATAC-seq time point with shuffled training, validation, and test sets (three-fold validation). The stability of the performance metrics as well as the stability of the returned downstream motif grammar was compared to the original optimized model training event (fold 1). All ChromBPNet models were implemented and trained using Keras (v2.5.0^148^), TensorFlow2 backend (v.2.5.1^149^), and the Adam optimizer^150^. Training was performed using a NVIDIA^®^ TITAN RTX GPU with CUDA v11.0 and cuDNN v8.3.0 drivers.

### ChromBPNet contribution score generation and validation

DeepLIFT (v0.6.13.0, derived from the Kundaje Lab fork of DeepSHAP (https://github.com/AvantiShri/shap)^151^) was applied to the trained ChromBPNet model to generate the contribution of each base across a given input sequence to the predicted output counts and profile signals. Contribution scores for counts and profile outputs were generated for each trained ChromBPNet model across all time points (1-1.5 h, 1.5-2 h, 2-2.5 h, 2.5-3 h). TF-MoDISco (v.0.5.16.0^152^) was then applied for each trained ChromBPNet model in order to identify regions of high counts contribution, cluster based on within-group contribution and sequence similarity, and consolidate these clusters into motifs. Pertinent motifs (Zelda, GAF, Caudal, Twist-like, and Dorsal-like) were manually identified based on similarity to previous literature and ChIP-nexus binding was measured across these accessibility-identified motifs to validate that they were indeed relevant binding sites that also contribute towards explaining the ChromBPNet models across the designated time points (Supplemental figure 6f).

### Using binding and accessibility models to examine motif effects *in silico*

In order to internally measure the “marginalized” effects of motifs without the surrounding genomic context, we adopted an *in silico* approach by which we injected motifs into many seed-controlled randomized sequences and generated BPNet and ChromBPNet predictions of these sequences with and without the motifs. We used 64 randomized sequences for BPNet predictions and 512 for ChromBPNet predictions (accessibility predictions contain greater sequence complexity and therefore required more trials to establish stable predictions across randomly generated sequences), averaging predictions across each of these randomized sequence sets. After performing *in silico* injections of a single motif, we visualized the output profiles generated from randomized sequence alone or motif-injected sequences for the Tn5 bias model, ChromBPNet models, and BPNet across all TF motifs.

It has been previously described that accurate predictions of relative motif affinities can be extracted from a BPNet model trained on ChIP-nexus data^92,93^. We then summarized the “marginalized” effects of motifs above to compare how motif affinity changes Zelda’s influence at the level of both binding and accessibility. After performing *in silico* injections of a single motif described above, we summed the values of the output profiles generated from randomized sequence alone or motif-injected sequences for both ChromBPNet and BPNet. These sums were then subtracted in log-space and referred to as “marginalized” scores, characterized as:

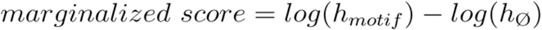

where *h_motif_* is the predicted sum of the counts when a motif is injected into the random sequence and *h_ø_* is the predicted sum of the counts of the averaged random sequences without injections. These “marginalized” scores were computed for each Zelda motif variant for all ChromBPNet models and BPNet.

In order to test the effects of motif pairs on cooperativity for binding and accessibility without surrounding genomic context, *in silico* motif interaction analysis was performed as described previously^57^. In brief, this involved injecting two motif sequences (motif A and motif B) across motif pair distances (*d*) ranging up to 400 bp into random sequences.

Binding predictions and accessibility predictions were measured in these different simulation scenarios from BPNet (where *h* represents the sum of the counts predicted across a 200 bp window, centered on motif A) and ChromBPNet (where *h* represents the sum of the counts predicted across the entire 1000 bp window), respectively. We measured four different cases: (1) neither motif A nor motif B were injected into the sequence (*hØ*), (2) motif A only was injected into the sequence *(hA)*, (3) motif B only was injected into the sequence *(hB)*, and (4) motif A and motif B were both injected into the sequence at a designated distance (*hAB*). These cases were measured and averaged across 64 trials for BPNet predictions and 512 trials for ChromBPNet predictions (accessibility predictions contain greater sequence complexity and therefore required more trials to establish stable predictions across randomly generated sequences). After all measurements were collected across all motif combinations and distances, then averaged across trials, the *in silico* motif pair cooperativity for each was calculated using the following equation:

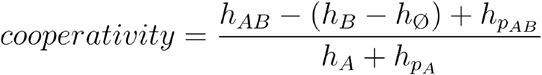

where (*h_p_*) is the predicted pseudocounts represented by the 20th percentile quantile cutoff value for both binding and accessibility predictions across each window when motif A and motif B are present and when only motif A is present (case 4 and 2, respectively, described above). The motif pairs considered were combinations of the highest affinity representations of Zelda (CAGGTAG), Dorsal (GGGAAAACCC), Twist (AACACATGTT), Caudal (TTTTATGGCC), Bicoid (TTAATCC), and GAF (GAGAGAGAGAGAGAGAG). For both BPNet and all ChromBPNet models, these high-affinity motifs were also tested alongside an additional lower affinity representation of Zelda (TAGGTAG) in a pairwise fashion with all other motifs to investigate Zelda’s changing influence on other TFs based on motif affinity.

### Using binding and accessibility models to examine motif effects in genomic sequences

In order to measure the in-context effects of a motif within its surrounding genomic sequence, we computationally generated genomic sequences with this motif’s sequence mutated by randomly shuffling the bases that belong to this motif. We generated 16 randomized mutation sequences per motif instance to establish mutation stability, averaging predictions across each of these randomized mutation sets. We performed this genomic perturbation for all mapped TF motifs across our curated set of genomic enhancers (described above) and visualized the output profiles generated for both BPNet and all ChromBPNet models.

In order to summarize the accessibility effects of mutating high- and low-affinity Zelda motifs, the 250 highest- and lowest-affinity Zelda motif-containing-only islands were identified. Using the procedure described above for all Zelda motifs in these genomic islands, accessibility profiles from unmodified island sequences and Zelda-mutated island sequences were predicted using the ChromBPNet models. After generating the profiles for each island, we summed the profiles into a single scalar value for WT sequences (*h_WT_*) and Zelda-mutated sequences (*h_dZld_*). Relative accessibility effects of high- and low-affinity Zelda motifs were characterized by the log2 fold-change measured effect, represented as 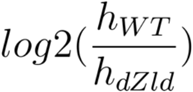.

### Differential chromatin accessibility analysis

To determine the differential chromatin accessibility between *wt* embryos with mutant *zld^-^, gd^7^*, and *cic^6^* embryos, we used DESeq2 with default parameters and FDR = 0.05^103^. Briefly, for each comparison between *wt* and mutant ATAC-seq data sets, we calculated ATAC-seq cut site coverage at the same pooled IDR-reproducible ATAC-seq peaks from all time points that were used for ChromBPNet prior to promoter removal (see “ChromBPNet model training and optimization”). For all time points we used three replicates and built one DESeq model encompassing ATAC-seq counts from all time points. In order to compute the differential chromatin accessibility, we then used each DESeq2 model to conduct pairwise comparisons between between *wt* and mutant conditions within each time point and computed the log2(mutant/*wt*) values. In this way, log2(mutant/*wt*) < 0 represent a loss in chromatin accessibility in the mutant, while log2(mutant/*wt*) > 0 represent a gain in chromatin accessibility in the mutant, while p-adjusted < 0.05 loci are highlighted. We performed this differential chromatin accessibility approach for all *wt*-to-mutant comparisons.

### Enhancer collection

The bulk set of mesodermal and dorsal ectodermal enhancers used in this study were previously defined based on differential histone acetylation^102^. More limited sets of validated neuroectodermal enhancers, as well as mesoderm and dorsal ectoderm enhancers, were collected from previous work^77,153^. All anterior-posterior patterning enhancers were collected from earlier studies^74,75^. Additional enhancer lists that were consulted include a list of active blastoderm enhancers^73^ and REDfly^154^.

## Supplementary figures

**Supplemental figure 1.**
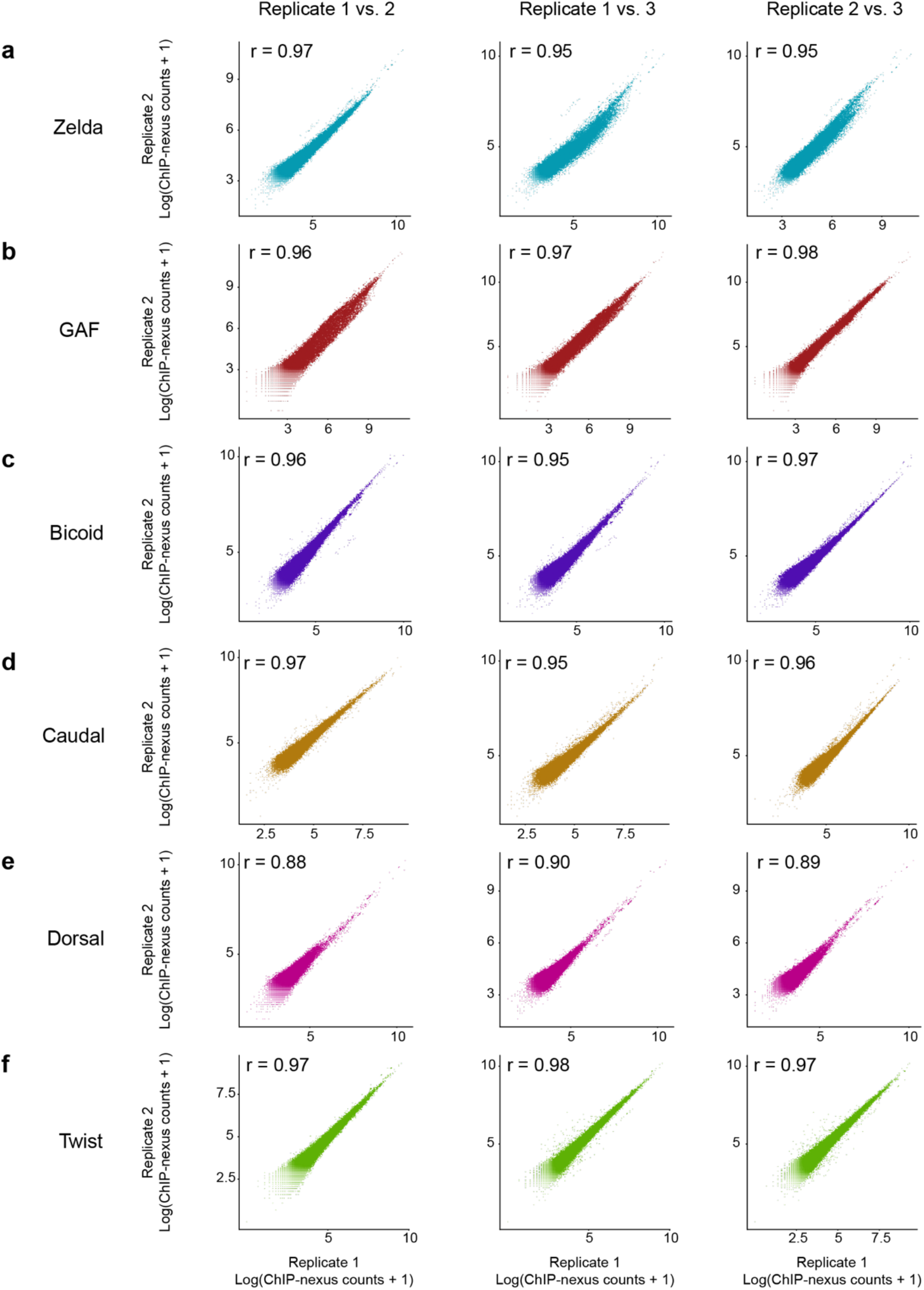
ChIP-nexus replicates for all TFs are highly correlated. Pearson correlation values were determined for the three replicates of **(a)** Zelda, **(b)** GAF, **(c)** Bicoid, **(d)** Caudal, **(e)** Dorsal, and **(f)** Twist ChIP-nexus experiments. Coverage for each replicate was calculated across a 400 bp window centered on the MACS2-called peaks for each TF. Because ChIP-nexus provides strand-specific information, the absolute value of the counts from the negative strand, which would otherwise be negative, was taken and added to the counts across the positive strand to determine the total region counts for a given replicate.

**Supplemental figure 2.**
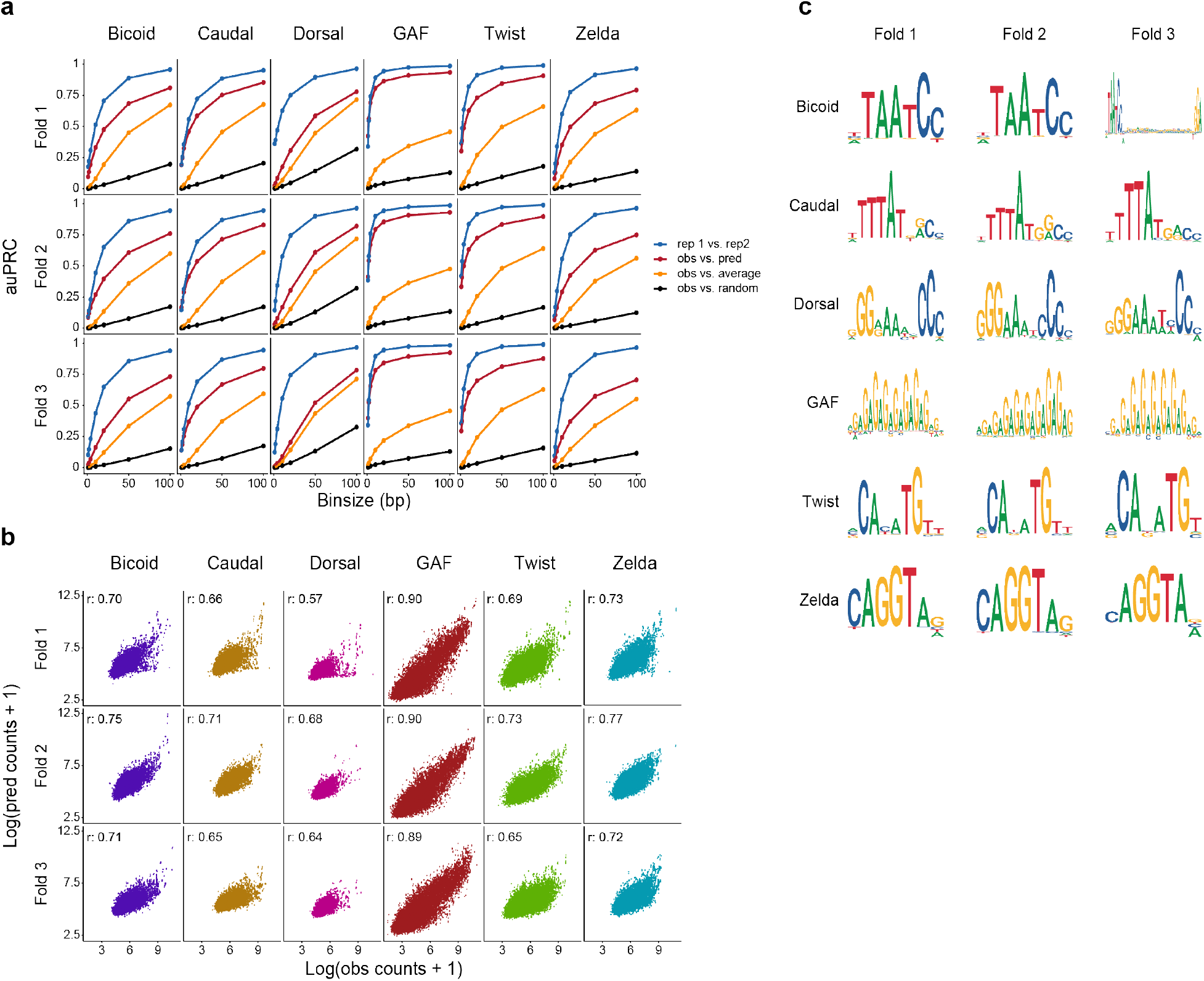
BPNet accurately learns the profile and counts information for all TFs of interest irrespective of the training chromosome set. **(a)** Area under the Precision-Recall Curves (auPRC) show that BPNet predicts the profile positions with high accuracy. The ability of BPNet to identify positions of high ChIP-nexus signal is assessed at various resolutions up to 100 bp. Replicate experiments, average ChIP-nexus profiles, and randomized profiles are shown as controls. Three-fold validation was performed by applying the same model architecture from the original, optimized model (fold 1) to two additional models (fold 2 and fold 3) with the training, validation, and test chromosomes shuffled. These results show that the training regions are representative of the entire dataset and that the trained BPNet model is highly stable. **(b)** BPNet predicts ChIP-nexus counts with high accuracy. Pearson counts correlation values were determined by comparing the observed ChIP-nexus counts with BPNet’s predicted counts at ChIP-nexus peaks for each of the TFs of interest. The stability of BPNet’s counts predictions were assessed with three-fold validation. **(c)** BPNet re-discovered the known motifs for all TFs of interest irrespective of the distribution of the training, validation, and test chromosomes. BPNet CWMs are shown for each TF for the original optimized model (fold 1) and the additional models trained with the same architecture as part of three-fold validation (fold 2 and fold 3).

**Supplemental figure 3.**
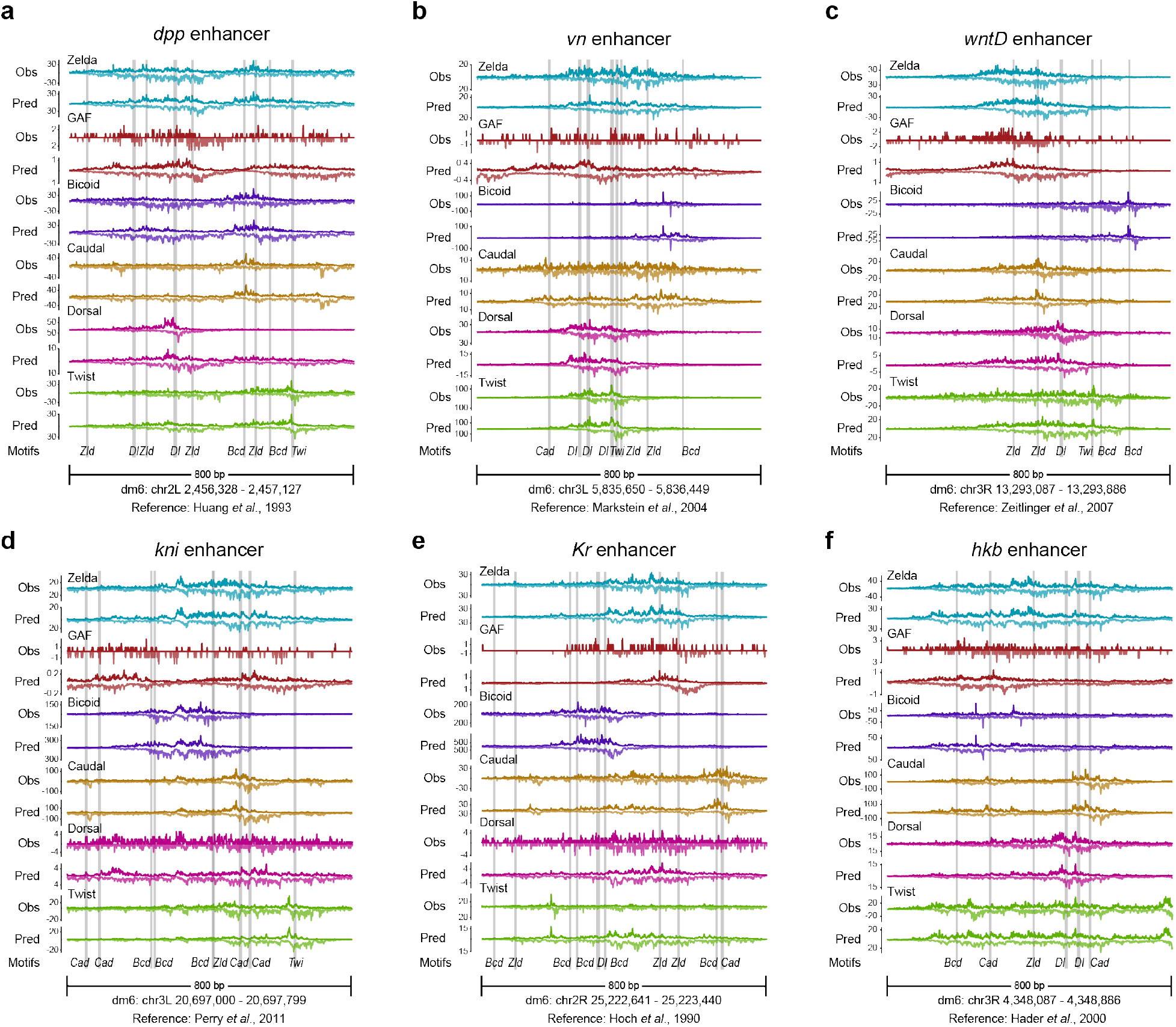
BPNet accurately maps TF motifs and predicts TF binding at known *Drosophila* enhancers. As in Figure 1d, the experimentally generated ChIP-nexus (top track) and BPNet predicted ChIP-nexus data (bottom track) for each TF (different colors) are plotted at known enhancers for the following genes: **(a)** *dpp*^155^, **(b)** *vn^156^*, **(c)** *wntD*^153^, **(d)** *kni*^157^, **(e)** *Kr*^158^, and **(f)** *hkb*^159^. Motifs were discovered and mapped by BPNet and references for each enhancer have been included. Enhancers across different developmental patterns and axes were deliberately selected for showcasing BPNet’s predictive accuracy.

**Supplemental figure 4.**
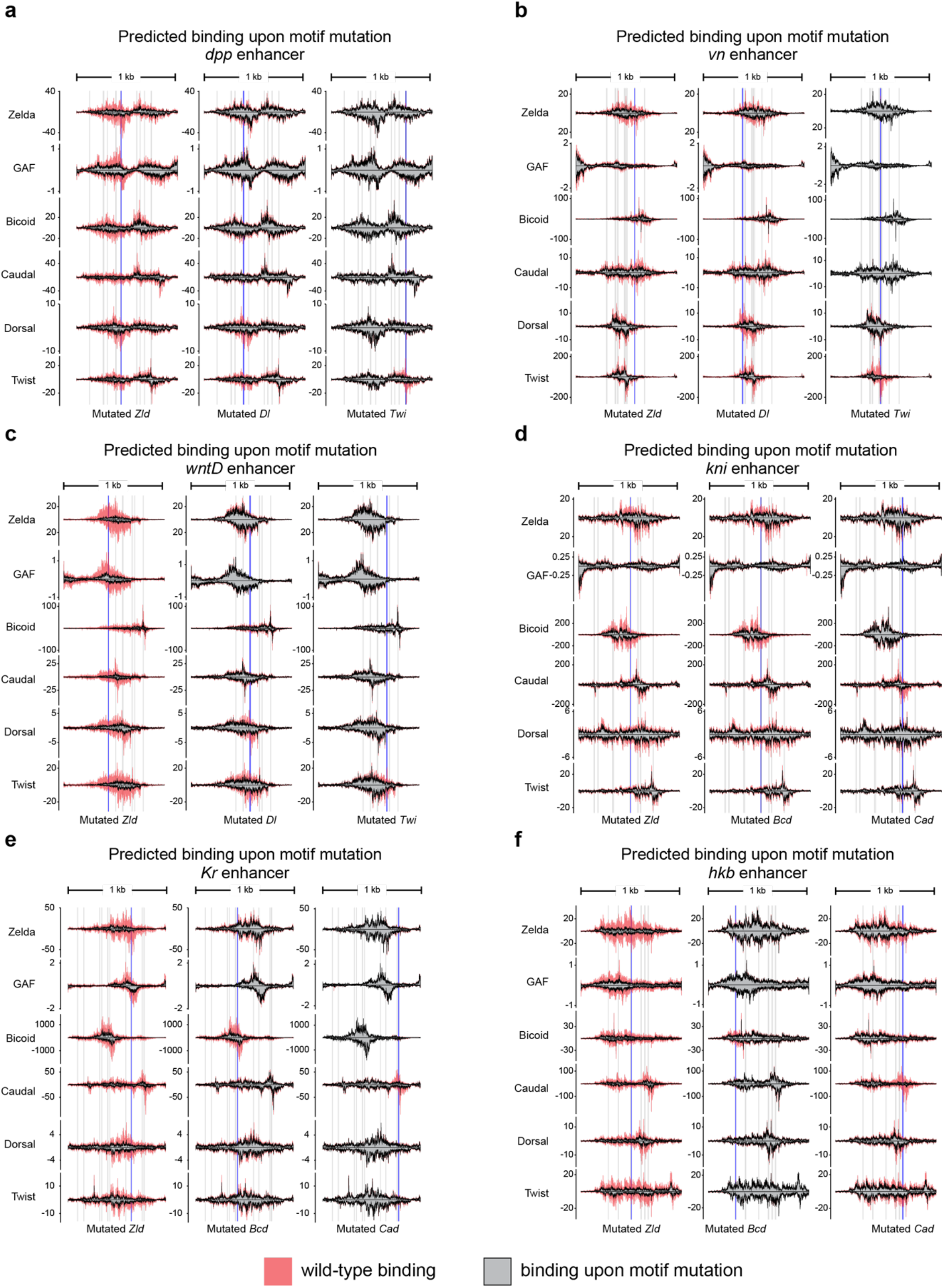
BPNet predicts the effects of single motif mutations at known *Drosophila* enhancers. As in Figure 1g, BPNet predicted the binding of all TFs at *wt* enhancer sequences and again at enhancers upon individual motif mutations for the following enhancers: **(a)** *dpp*, **(b)** *vn*, **(c)** *wntD*, **(d)** *kni*, **(e)** *Kr*, and **(f)** *hkb*. Shaded colors show TF binding across the *wt* enhancer, while the gray-filled profiles represent TF binding in response to the motif mutation. Blue bars indicate the mutated motifs that are highlighted under the predictions, and the gray bars are all other BPNet-mapped motifs across the enhancers. Enhancer coordinates and references are provided in Supplemental figure 3.

**Supplemental figure 5.**
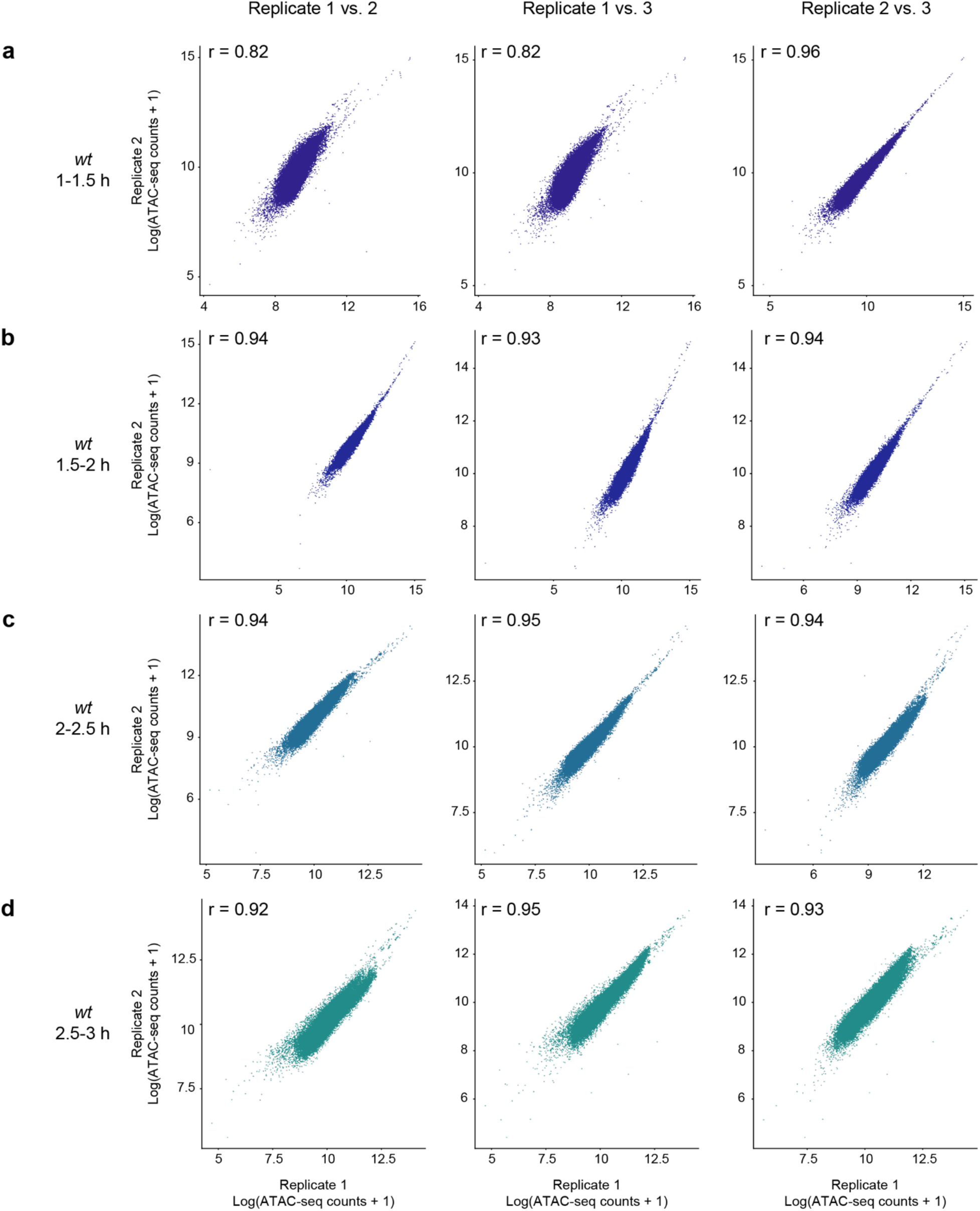
Time course ATAC-seq experiments in *wt* embryos are highly correlated. Pearson correlation values were determined for the three replicates of **(a)** 1-1.5 h AEL, **(b)** 1.5-2 h AEL, **(c)** 2-2.5 h AEL, and **(d)** 2.5-3 h AEL *wt* ATAC-seq experiments. ATAC-seq counts for each replicate were calculated across a 400 bp window centered on the MACS2-called peaks for each time point.

**Supplemental figure 6.**
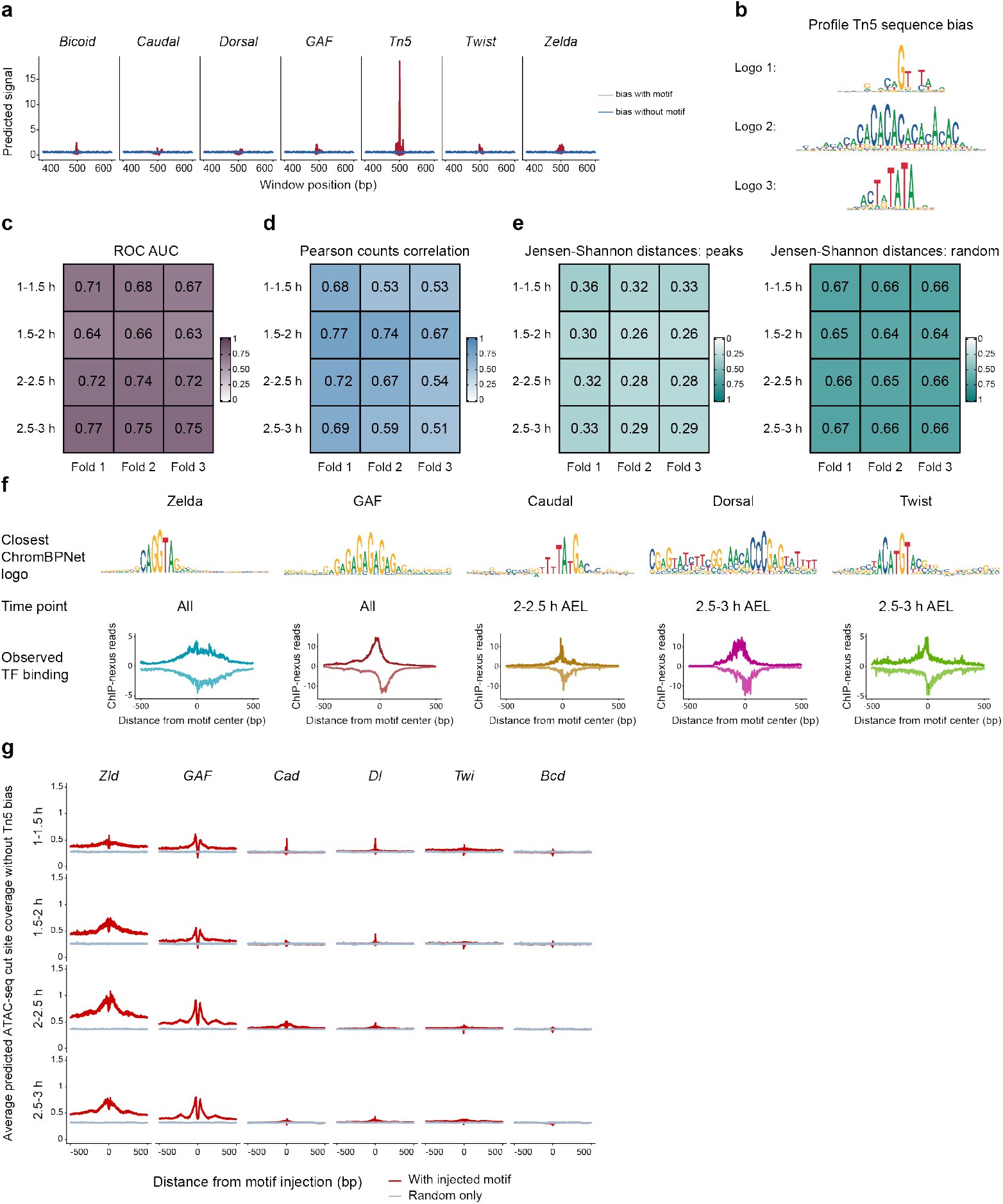
ChromBPNet accurately learns time course chromatin accessibility without Tn5 bias in the early *Drosophila* embryo. **(a)** The Tn5 bias ChromBPNet model does not learn TF motif sequence grammar. The canonical sequences for each TF of interest were injected into 256 genomic sequences from ChromBPNet’s test chromosome (chrX) and the Tn5 bias model was used to predict chromatin accessibility cut site signal. The effects were averaged across trials and show no predicted accessibility upon injection of any motif except the Tn5 preferred sequence. This confirms that the bias model’s learning was limited to Tn5 bias and did not learn cis-regulatory grammar. **(b)** The Tn5 sequence bias is represented by multiple sequence logos. TF-MoDISco interpretations returned Tn5 sequence bias as multiple logos for profile contribution but not for counts contribution. These results show that the bias model only learned Tn5 positional information and was successfully trained to only represent Tn5 bias at closed genomic regions. **(c)** The time course ChromBPNet models accurately discriminates between ATAC-seq peak and nonpeak regions. The models’ predictions were assessed using area under the receiver operating characteristic curves (ROC AUC). Three-fold validation was performed as in Supplemental figure 2 by applying the original ChromBPNet architecture (fold 1) to two additional models with reshuffled training, test, and validation chromosome sets (fold 2 and fold 3). **(d)** The time course ChromBPNet models accurately predict chromatin accessibility counts. Pearson correlation values were calculated by comparing the observed ATAC-seq cut sites with the ChromBPNet predicted cut sites at ATAC-seq peak regions for all time points. Three-fold validation was performed as described above. **(e)** The time course ChromBPNet models have high profile prediction accuracy. Time course profile predictions were assessed by comparing to the observed ATAC-seq cut sites using Jensen-Shannon distances at peak regions, where lower values are better. Randomly shuffled region profiles were included as a control. Three-fold validation was performed as described above. **(f)** ChromBPNet identifies TF motifs in ATAC-seq data that are bound by their respective TFs. TF-MoDISco was run on all ChromBPNet models and sequence features with high counts contribution were consolidated into motifs. The closest motif logo for each TF of interest was manually identified with the exception of Bicoid, which ChromBPNet did not identify in the ATAC-seq data. Motifs for the pioneering TFs were unambiguous and identified at all time points, while the patterning TF motifs deviated from the BPNet-identified binding motifs for Caudal, Dorsal, and Twist and were identified only at later time points. Average observed ChIP-nexus binding profiles showed clear TF footprints on all motifs. Average footprints were anchored on and calculated across the accessibility-identified motifs for each TF. **(g)** ChromBPNet predicts time course chromatin accessibility in response to TF motif injection. The TF binding motifs identified by BPNet were injected into 512 randomized sequences. The ChromBPNet models were used to make chromatin accessibility cut site predictions before (blue) and after (red) TF motif injection, with the predicted effect centered on the injected motif. The predictions were averaged for all trials and show that pioneering motifs have the largest predicted effect on chromatin accessibility.

**Supplemental figure 7.**
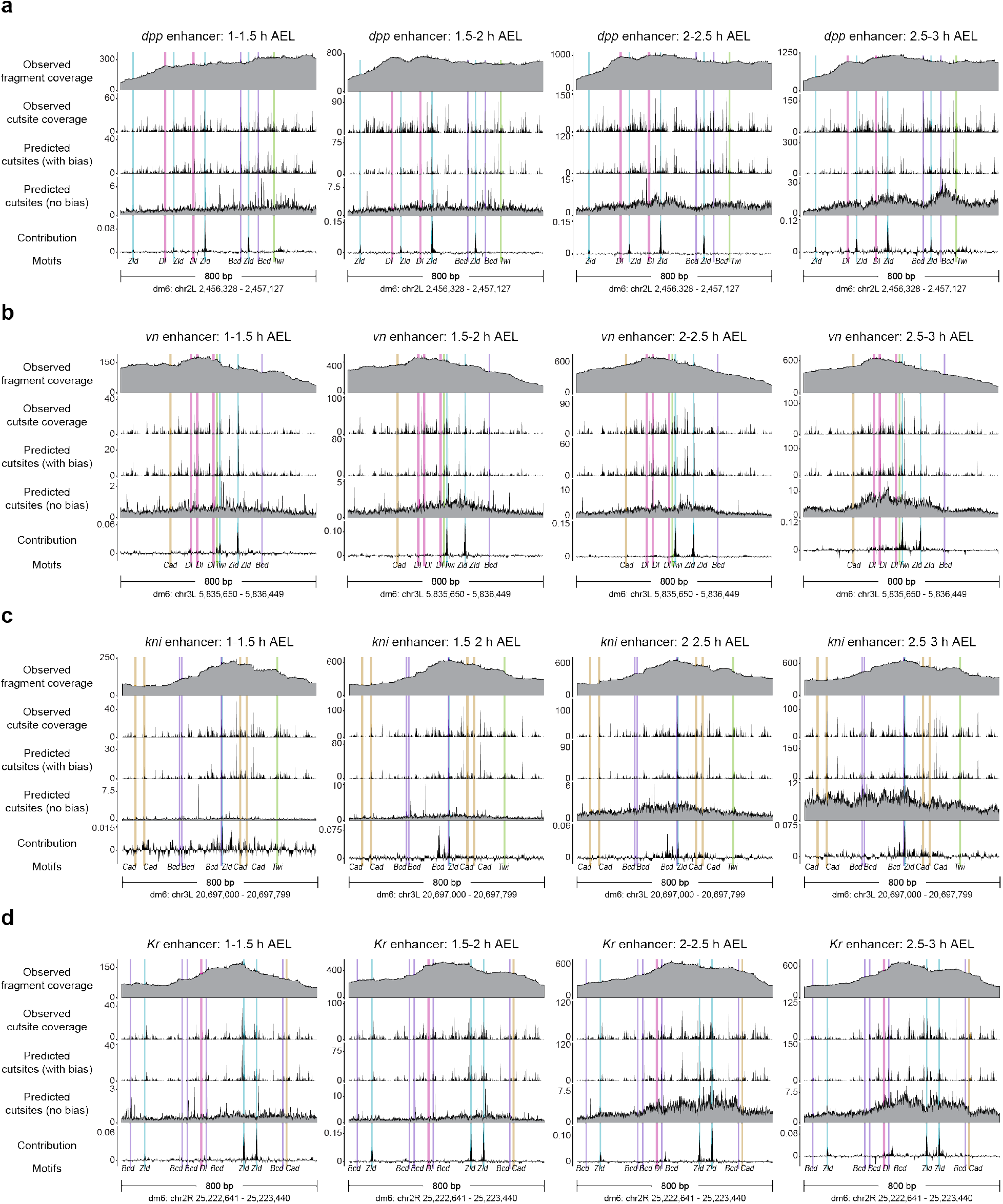
ChromBPNet predicts time course chromatin accessibility at known enhancers and identifies the TF motif contribution to accessibility. As in Figure 2c, the experimentally generated ATAC-seq data (tracks one and two) are shown with ChromBPNet accessibility predictions with Tn5 bias (track three) and without Tn5 bias (track four) for known enhancers for the following genes: **(a)** *dpp*, **(b)** *vn*, **(c)** *kni*, and **(d)** *Kr*. Columns represent model predictions at each of the four ATAC-seq time points. The counts contribution for chromatin accessibility across each enhancer is shown as the fifth track, with spikes at BPNet-mapped TF motifs. TF motifs are highlighted with the colored bars.

**Supplemental figure 8.**
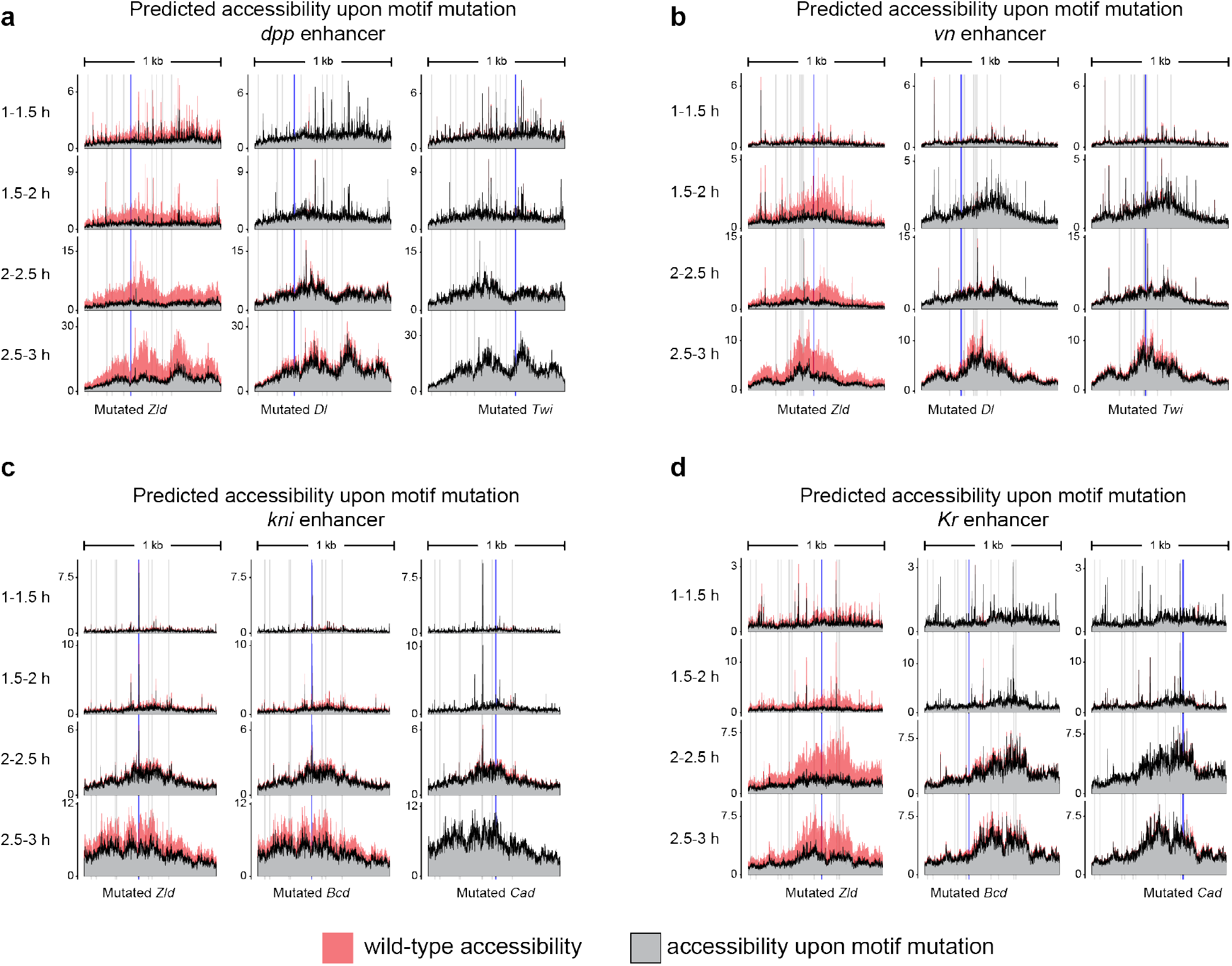
ChromBPNet predicts the effects of single motif mutations at known *Drosophila* enhancers. As in Figure 2d, ChromBPNet predicted chromatin accessibility cut sites without Tn5 bias for all time points at *wt* enhancer sequences and when individual motifs are mutated. This is performed at enhancers for the following genes: **(a)** *dpp*, **(b)** *vn*, **(c)** *kni*, and **(d)** *Kr*. Shaded colors show chromatin accessibility predictions across the *wt* enhancer sequence, and the gray-filled profiles represent chromatin accessibility upon mutation of the highlighted motif (blue bar). Gray bars are non-mutated motifs across each enhancer.

**Supplemental figure 9.**
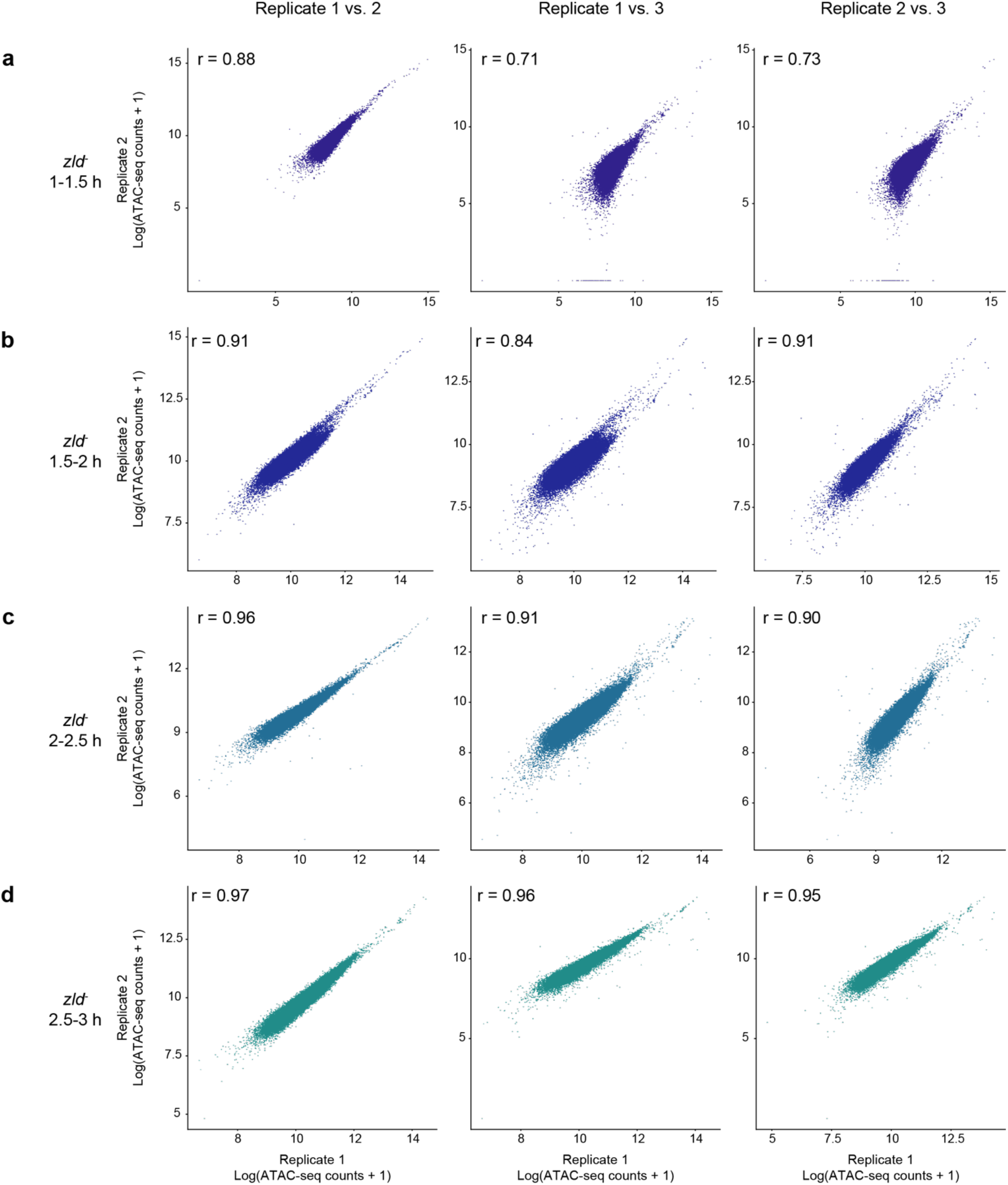
Time course ATAC-seq experiments in *zld^-^* embryos are highly correlated. Pearson correlation values were determined for the three replicates of **(a)**1-1.5 h AEL, **(b)**1.5-2 h AEL, **(c)**2-2.5 h AEL, and **(d)**2.5-3 h AEL *zld^-^* ATAC-seq experiments. ATAC-seq counts for each replicate were calculated across a 400 bp window centered on the MACS2-called peaks for each time point.

**Supplemental figure 10.**
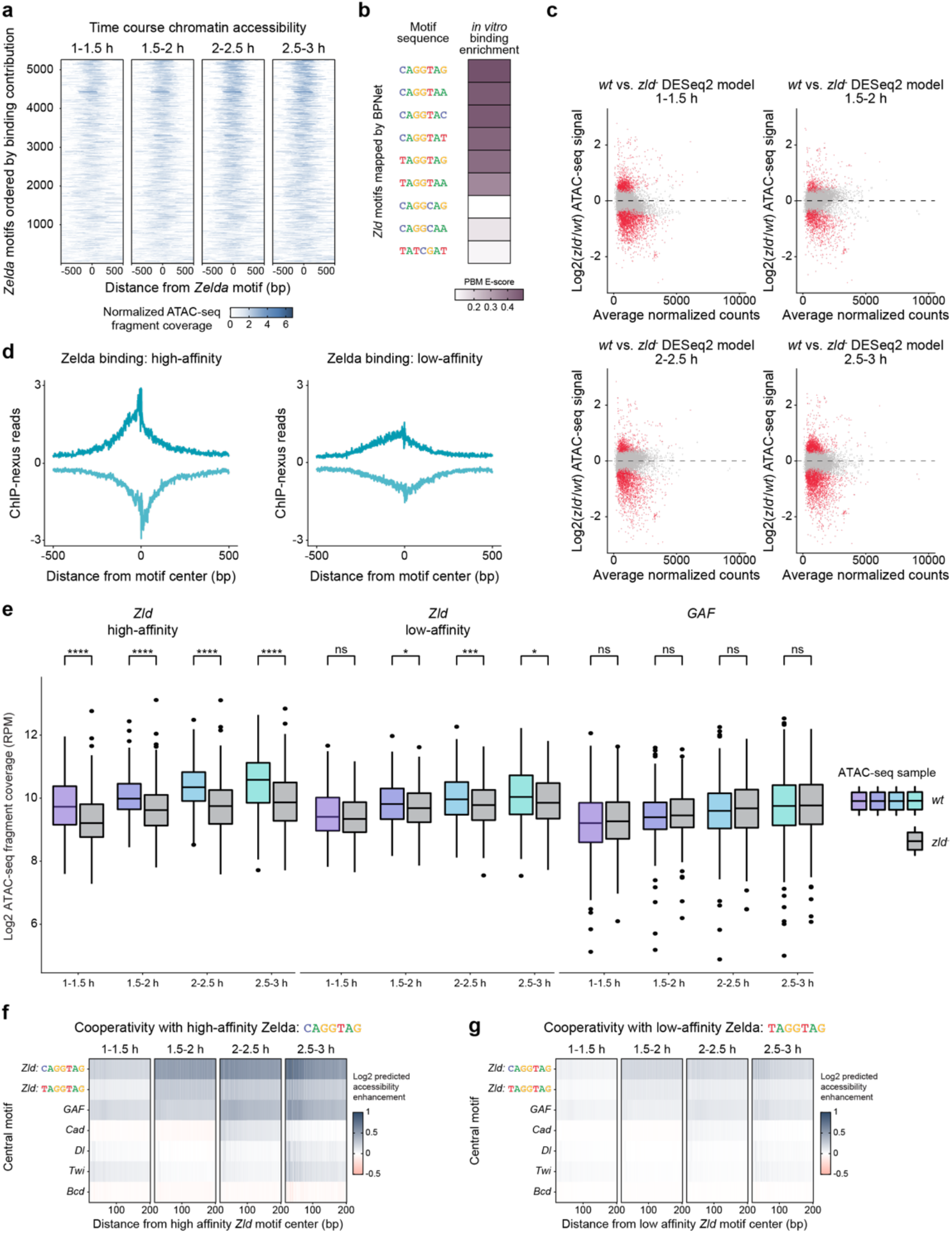
High-affinity Zelda motifs have greater effects on chromatin accessibility than low affinity Zelda motifs. **(a)** Time course chromatin accessibility correlates with Zelda motif binding contribution. BPNet-mapped Zelda motifs were ordered by their counts contribution scores for Zelda binding as in Figure 3a. The experimentally generated ATAC-seq signal was calculated across a 1000 bp window, anchored on the Zelda motif, for each time point. The Zelda motifs that contribute most strongly to Zelda binding exhibit the highest chromatin accessibility. **(b)** Protein binding microarray (PBM) E-scores show differences between high- and low-affinity Zelda motifs. The Escore is a rank-based PBM statistic that is a variation on the area under the receiver operating characteristic curve (AUC) that ranges from −0.5 (lowest) to 0.5 (highest)^87^. E-scores for each Zelda heptad were calculated as done in Figure 3b. **(c)** Time course MA plots show differential chromatin accessibility between *wt* and *zld^-^* embryos. The differential chromatin accessibility was calculated between *wt* and *zld^-^* embryos using DESeq2 for all time points. Red highlighted dots are ATAC-seq peaks that are differentially accessible with statistical significance (FDR = 0.05). **(d)** Zelda is more strongly bound to high-affinity motifs than to low-affinity motifs. Average Zelda binding footprints were calculated and plotted across the same high- and low-affinity Zelda motifs as in Figure 3f. Average profiles were calculated across a 1000 bp window and were anchored on Zelda motifs. **(e)** Low-affinity Zelda motifs have a weaker effect on chromatin accessibility than high-affinity Zelda motifs. The average profiles in Figure 3f were quantified using boxplots and were tested for statistical significance using the using the Wilcoxon rank-sum test (* = p < 0.05; ** = p < 0.01; *** = p < 0.001; **** = p < 0.0001). Observed ATAC-seq fragment coverage was calculated across a 500 bp window centered on each Zelda motif using the same motif instances as in Figure 3f. There is an average of a five-fold weaker effect from low-affinity motifs than from high-affinity motifs, which was calculated using median values for accessibility for *wt* and *zld^-^* embryos at all time points. **(f, g)** ChromBPNet predicts that high-affinity Zelda motifs induce greater chromatin accessibility than low-affinity Zelda motifs. *In silico* motif injections into randomized sequences were performed as in Figure 3h, except the ChromBPNet models were used to predict chromatin accessibility at TF motifs upon injection of a **(f)** high-affinity or **(g)** low-affinity Zelda motif for each time point. Motif injections were repeated 512 times and predictions were averaged.

**Supplemental figure 11.**
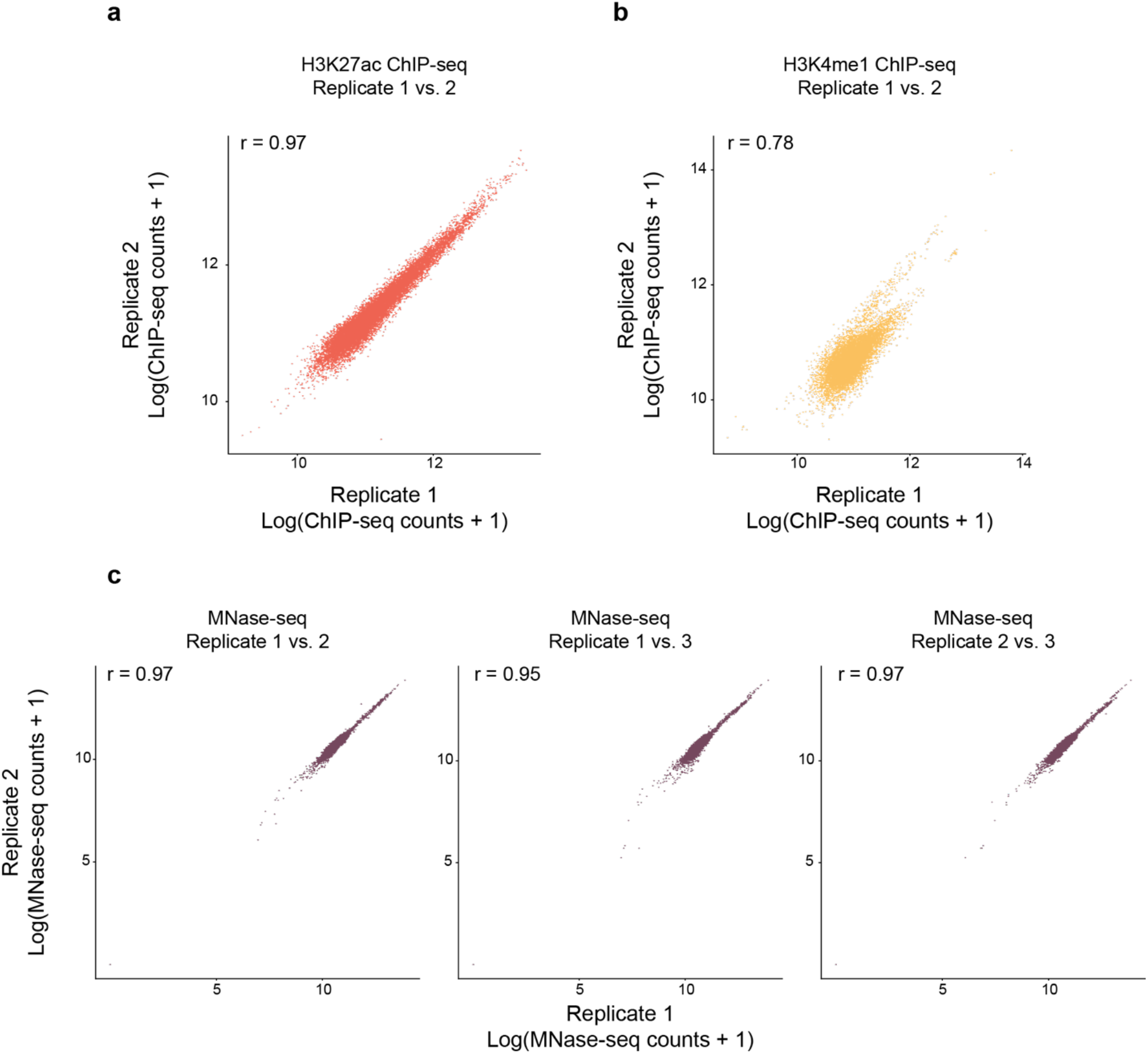
Histone modification ChIP-seq replicates and MNase-seq replicates are highly correlated. **(a, b)** Pearson correlation values were determined for the two replicates of **(a)** H3K27ac ChIP-seq and **(b)** H3K4me1 ChIP-seq. ChIP-seq counts for each replicate were calculated across a 1000 bp window centered on the MACS2-called peaks for each histone mark. **(c)** Pearson correlation values were determined for the three replicates of MNase-seq experiments. MNase-seq counts for each replicate were calculated across a 1000 bp window centered on *Drosophila* transcription start sites.

**Supplemental figure 12.**
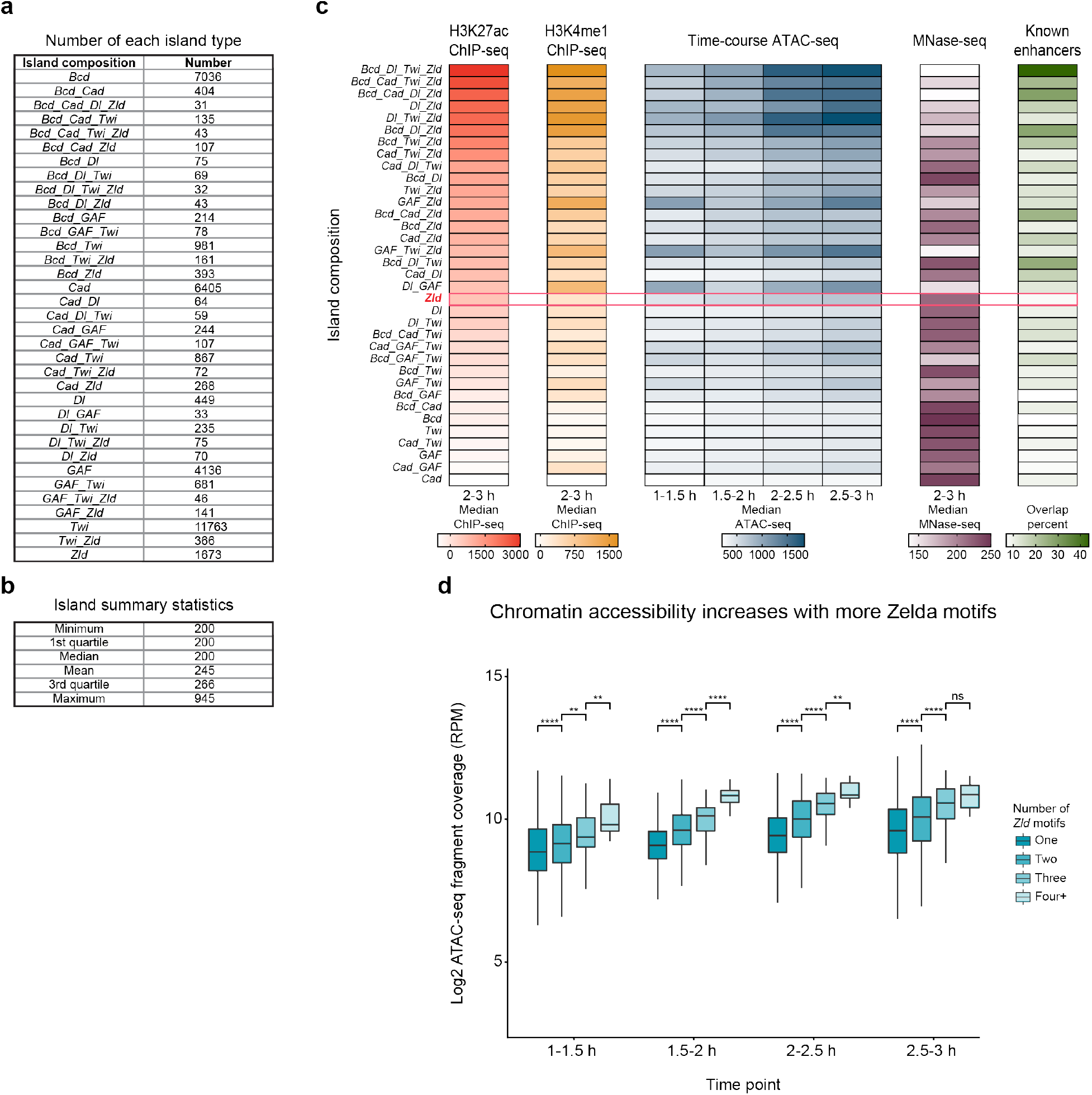
Chromatin accessibility correlates with the number of Zelda motifs in an island. **(a)** Summary of motif islands generated according to the scheme in Figure 4a. Only motifs identified and mapped by BPNet and that are bound by their associated TF are used for island generation. Islands with fewer than 30 genomic instances are excluded. Motif islands are not separated according to how many motifs they contain but are instead classified based on which TF motifs compose them. **(b)** Summary statistics for motif islands. Islands that are 200 bp wide are single-motif islands. Islands are approximately the sizes as *Drosophila* enhancers, with no island greater than 945 bp. **(c)** Motif islands of the same composition are grouped together and genomics signals for each island type are calculated exactly as in Figure 4b. Here, islands are ordered by H3K27ac signal. **(d)** Greater chromatin accessibility is associated with regions with more mapped Zelda motifs. All Zelda-containing islands were collected and separated based on how many Zelda motifs they contained. The observed normalized ATAC-seq fragment coverage for each time point was calculated across a 250 bp window anchored on the island center. Statistical significance was determined using the Wilcoxon rank-sum test (* = p <0.05; ** = p < 0.01; *** = p < 0.001; **** = p < 0.0001). These results show that more Zelda motifs across a genomic region correlates with increased chromatin accessibility. This is consistent with previous results showing higher levels of nucleosome depletion for more Zelda motifs^6^.

**Supplemental figure 13.**
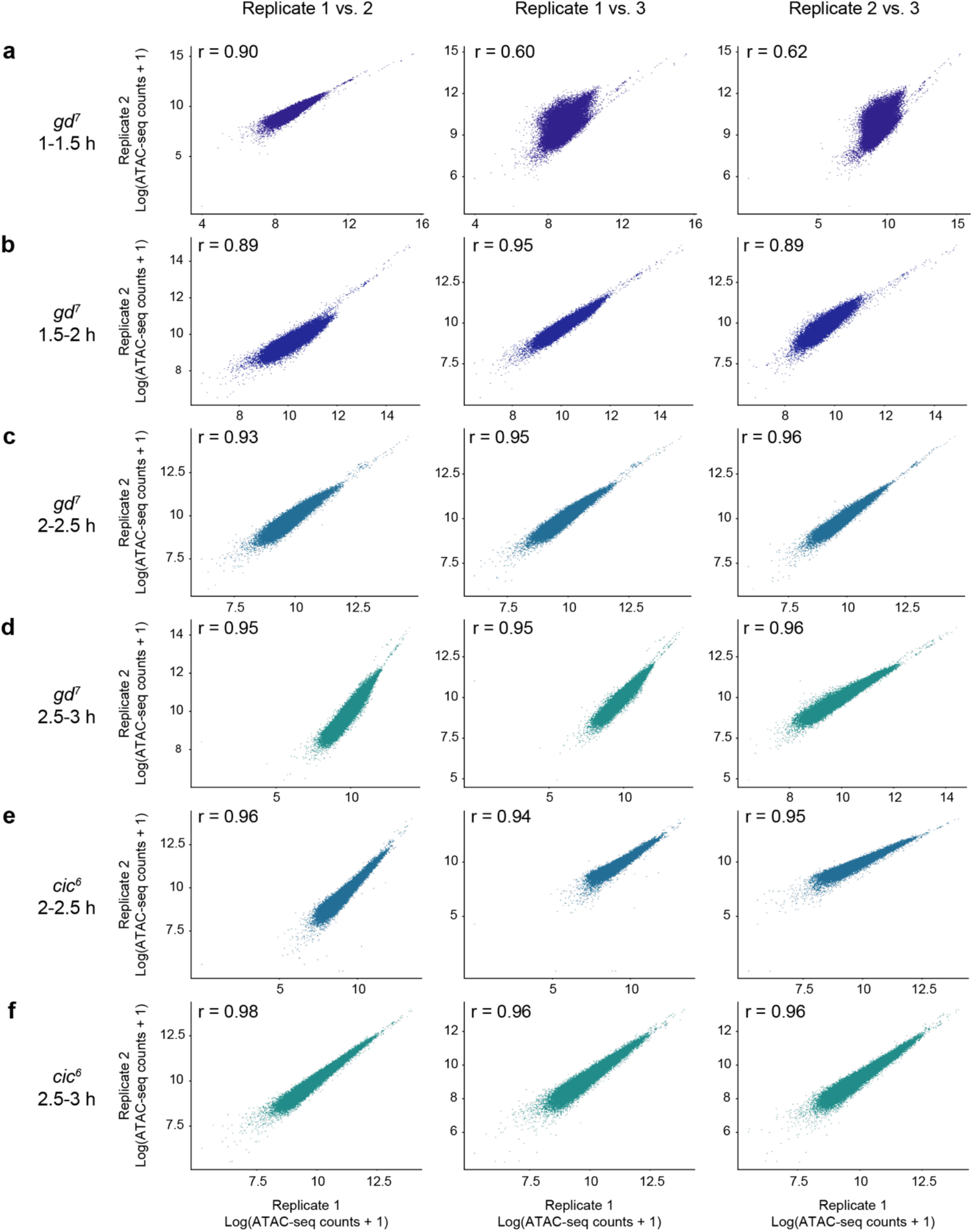
Time course ATAC-seq replicates are highly correlated in *gd^7^* and *cic^6^* embryos. **(a-d)** Pearson correlation values were determined for the three replicates of **(a)** 1-1.5 h AEL, **(b)** 1.5-2 h AEL, **(c)** 2-2.5 h AEL, and **(d)** 2.5-3 h AEL *gd^7^* ATAC-seq experiments. ATAC-seq counts for each replicate were calculated across a 400 bp window centered on the MACS2-called peaks for each time point. **(e, f)** Pearson correlation values were determined for the three replicates of **(e)** 2-2.5 h AEL and **(f)**2.5-3 h AEL *cic^6^* ATAC-seq experiments. ATAC-seq counts for each replicate were calculated across a 400 bp window centered on the MACS2-called peaks for each time point.

**Supplemental figure 14.**
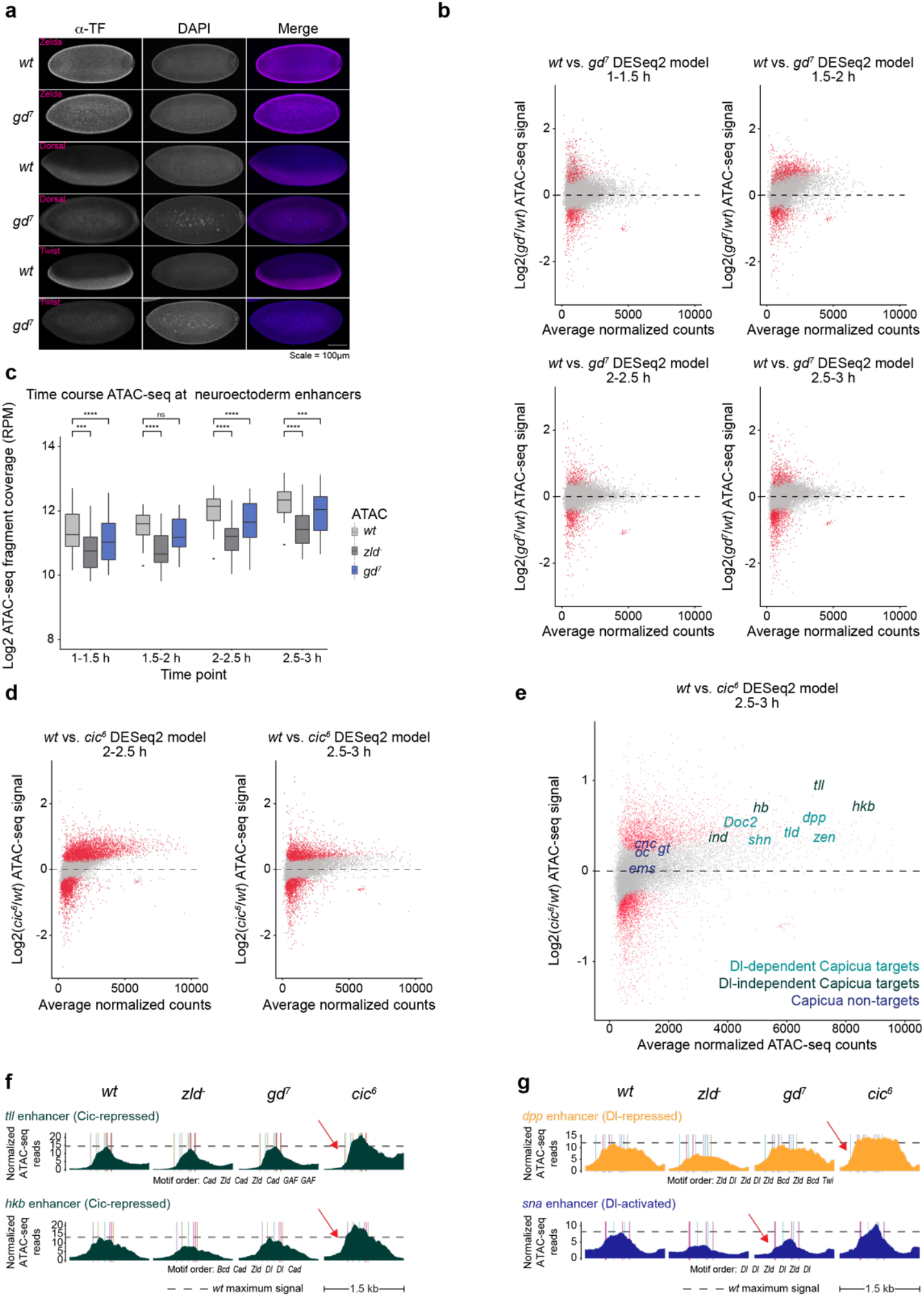
Chromatin accessibility changes context-specifically in *gd^7^* and *cic^6^* embryos. **(a)** *gd^7^* embryos show a clear loss of dorsoventral patterning. Nuclear cycle 14 embryos were stained using the same Zelda, Dorsal, and Twist antibodies used in ChIP-nexus experiments. Confocal images of *wt* and *gd^7^* embryos were collected using the same settings, maximum intensity projected, and processed in FIJI using the identical settings. **(b)** Time course MA plots show differential chromatin accessibility between *wt* and *gd^7^* embryos. DESeq2 was used to determine differential chromatin accessibility for all time points, and the red points represent the ATAC-seq peaks that are significantly differentially expressed (FDR = 0.05). **(c)** Neuroectoderm enhancers lose chromatin accessibility in *gd^7^* embryos. The normalized ATAC-seq fragment coverage was calculated in wt, *zld^-^*, and *gd^7^* embryos across known neuroectoderm enhancers (n = 23)^77,153^ as in Figures 5c and 5d. Wilcoxon ranksum tests were used to test for statistical significance (*** = p < 0.001; **** = p < 0.0001). In *gd^7^* embryos, neuroectoderm enhancers are inactive. **(d)** MA plots show differential chromatin accessibility between *wt* and *cic^6^* embryos at 2-2.5 h AEL and 2.5-3 h AEL. DESeq2 was run on *wt* and *cic^6^* embryos for both time points to determine the differential chromatin accessibility. ATAC-seq peaks that are significantly differentially accessible are highlighted in red (FDR = 0.05). **(e)** Chromatin accessibility is increased at Dorsal-independent Capicua-repressed enhancers in *cic^6^* embryos. Capicua represses known anterior-posterior enhancers (e.g., *hb, tll, hkb)* and the neuroectoderm enhancer *ind* without requiring Dorsal binding. Differential chromatin accessibility analysis was performed between *wt* and *cic^6^* embryos as in Figure 5e. Both Dorsal-dependent (teal) and Dorsal-independent (green) enhancers gain accessibility in *cic^6^* embryos, while enhancers not bound by Capicua (blue) do not. **(f)** Summary of chromatin accessibility at two Dorsal-independent Capicua-repressed enhancers (*tll* and *hkb*) upon loss of Zelda, nuclear Dorsal, and Dorsal-mediated repression as in Figure 5f. The dm6 enhancer coordinates are chr3R 30,851,400 - 30,852,900 (*tll*) and chr3R 4,347,821 - 4,349,321 *(hkb)*. Both enhancers do not significantly change accessibility in *gd^7^* embryos but do show increased accessibility in *cic^6^* as they are de-repressed (red arrows show DESeq2 statistical significance). **(g)** Summary of chromatin accessibility at a Dorsal-repressed enhancer *(dpp)* and Dorsal-activated enhancer *(sna)* upon loss of Zelda, nuclear Dorsal, and Dorsal-mediated repression as in figure 5f. The dm6 enhancer coordinates are chr2L 2,456,160 - 2,457,660 *(dpp)* and chr2L 15,479,300 - 15,480,800 (*sna*).

